# Comparative assessment of genomic, phenomic, and metabolomic prediction models in biparental grapevine breeding populations

**DOI:** 10.1101/2025.10.24.684307

**Authors:** Clémentine Borrelli, Louise Delannoy, Hadrien Chepca, Marcos Calcaterra, Elsa Chedid, Guillaume Arnold, Vincent Dumas, Raymonde Baltenweck, Alessandra Maia-Grondard, Philippe Hugueney, Didier Merdinoglu, Éric Duchêne, Komlan Avia

**Affiliations:** INRAE, Université de Strasbourg, SVQV, 68000 Colmar, France; Département de Phytologie, Faculté des sciences de l’agriculture et de l’alimentation, Université Laval, Québec City, Québec, Canada; Centre de recherche et d’innovation sur les végétaux, Université Laval, Québec City, Québec, Canada; Institut de Biologie Intégrative et des Systèmes, Université Laval, Québec City, Québec, Canada; EGFV, Univ. Bordeaux, Bordeaux Sciences Agro, INRAE, ISVV, 33882 Villenave d’Ornon, France

## Abstract

Accelerating grapevine breeding for disease resistance and climate adaptation remains constrained by long generation cycles. We benchmarked genomic (SNP), phenomic (NIRS), and metabolomic (untargeted LC-MS) prediction for 24 agronomic traits in a biparental population phenotyped over three years. Seven statistical frameworks and four tissue x timepoint combinations (wood; vineyard leaves at budbreak and flowering; greenhouse leaves at flowering) were evaluated, together with feature-wise BLUPs across samples. Cross-year and cross-population analyses with two additional populations assessed temporal robustness and transferability. Genomic prediction was most accurate (up to r = 0.83), metabolomic prediction was intermediate (up to r = 0.59), and phenomic prediction was lowest (up to r = 0.39) despite its lower acquisition cost. Metabolite features were more heritable than NIR wavelengths, for which most unexplained variation remained residual under the fitted model. Multi-omics integration produced limited overall gains. These results support genomic selection as the primary approach, with metabolomic or phenomic screening considered only for traits and sampling designs that show reproducible predictive signal.

## Introduction

Grapevine (*Vitis vinifera* L.) underpins high-value chains for wine, fresh fruit, and dried grapes. In 2025, the world vineyard covered ∼7.0 million hectares, and recent OIV (International Organization of Vine and Wine) assessments indicate that roughly half of the global fresh grape output is pressed (wine, musts, juices) while the remainder is consumed as table grapes or dried, underscoring the crop’s diversified economic relevance^1^. In France, viticulture occupies about 3 % of the Utilized Agricultural Area (UAA), yet it concentrates a disproportionate share of crop-protection inputs^2^. Fungicides account for the vast majority of treatments in vineyards, and in line with the EU Green Deal/Farm-to- Fork strategy, policy targets include a 50 % reduction in the use and risk of chemical pesticides by 2030^3^. These pressures, combined with climate change (earlier phenology and altered berry composition^4^), are forcing a shift toward varieties that can deliver disease resistance and maintain quality under warmer, drier, and more variable conditions.

Breeding disease-resistant grapevine cultivars, primarily to downy mildew (*Plasmopara viticola*), powdery mildew (*Erysiphe necator*), and black rot (*Phyllosticta ampelicida* a.k.a *Guignardia bidwellii*), typically requires introgression of resistance loci from North American or Asian *Vitis* species, while preserving agronomic performance and organoleptic quality^3,5^. European programs (e.g., JKI/WBI in Germany, INRAE-ResDur in France, FEM/VCR/University of Udine in Italy) have released disease resistant varieties (DRVs) and demonstrated substantial reductions in fungicide needs^6^, yet varietal development remains lengthy for a perennial crop. Shortening breeding cycles is therefore strategic for keeping pace with evolving pathogen populations and climate trajectories. While marker-assisted selection (MAS) is useful for pyramiding major resistance loci tightly linked to markers^7^, it is far less efficient for polygenic, quantitatively inherited traits (yield components, berry composition, phenology). Genomic selection (GS) addresses this limitation by fitting genome-wide markers to estimate genomic estimated breeding values (GEBV) and enable early selection^8^. Prediction accuracy depends on training population size and composition^9^, marker density and LD with causal variants^10^, genetic relatedness between training and target sets^11^, trait genetic architecture^12^, trait heritability^13^, and model choice^14^; optimizing training sets can substantially improve performance.

Beyond genotypes, endophenotypes (molecular or spectral fingerprints that aggregate genetic and environmental information) have emerged as powerful predictors^15^. Near-infrared spectroscopy (NIRS) enables “phenomic selection” (PS): using spectra to build relationship kernels and predict complex traits at very low cost and high throughput; PS can also model genotype-by-environment (GxE) patterns^16^. Likewise, metabolomic profiles (untargeted or targeted) have improved prediction in several crops, and multi-omics integration can further boost accuracy^17^. However, the cost and logistics of transcriptomic/metabolomic assays can be prohibitive in routine breeding, and their value relative to PS remains context-dependent.

In grapevine, GS can reach moderate-to-high predictive ability depending on trait architecture, linkage disequilibrium, and population structure^18,19^. Most prior work has focused on *V. vinifera* diversity panels or single-cross populations, with fewer evaluations in interspecific breeding backgrounds^18^. NIRS-based phenomic prediction (PP) has been evaluated at scale in grapevine and offers a low-cost complement to genomic prediction (GP)^19^. By contrast, metabolomic prediction (MP) has not, to our knowledge, been benchmarked alongside GP and PP across grapevine breeding populations, despite the broad use of metabolomics in grape and wine research^20^. Using a biparental population phenotyped over three years and genotyped at high density^21^, we (i) quantified feature-level genetic and residual variation in NIR wavelengths and metabolites; (ii) compared tissue and phenological-stage sampling strategies, including greenhouse-to-vineyard transfer; (iii) benchmarked seven models for GP, PP, and MP; (iv) evaluated multi-omics integration; and (v) assessed temporal and cross-population transferability. This design provides a side-by-side assessment of the accuracy, robustness, and practical limits of these omics-assisted breeding strategies.

## Results

The primary comparison comprised 24 agronomic traits and seven prediction frameworks. For matched cross-omics analyses, the 150 individuals selected for metabolomics were used for SNP and NIR prediction. The genomic matrix contained 7,386 SNPs, each NIR spectrum contained 2,332 wavelengths, and the LC-MS datasets contained 24,814 leaf features or 5,778 wood features. NIR and metabolomic predictors were evaluated for four sampling categories and as feature-wise BLUPs (genotype-level predictions for each wavelength or LC-MS feature after adjustment for sampling category).

### Effect of spectra pre-treatment

NIR spectra were acquired from wood and leaves collected at budbreak and flowering in the vineyard, and from leaves collected at flowering in the greenhouse. Spectra were analyzed either without transformation or after a Savitzky-Golay first derivative. Spectral profiles were broadly consistent among wood and vineyard leaf samples, although variability increased above 5,000 cm-1 and particularly above 7,000 cm-1; greenhouse leaves showed a more uniform variance profile. Wood spectra were less variable than leaf spectra. The first derivative reduced low-wavenumber baseline variability, whereas variability increased again at the upper end of the spectral range (Figure S1).

Because phenomic predictive abilities for all traits combined and several preprocessing-model combinations were close to zero and their standard deviations were often similar to or larger than their means (Table S1), preprocessing effects were interpreted cautiously. The first derivative had little effect on BayesC and RKHS predictions for greenhouse leaves at flowering, reduced the mean performance of LASSO, EN, and Random Forest, and slightly improved rrBLUP. For wood and vineyard leaves, it generally improved rrBLUP, BayesC, RKHS, and Random Forest predictions (Figure S2; Table S1), consistent with previous reports^19,22^. First-derivative spectra were therefore used in subsequent phenomic analyses, but the observed differences should not be interpreted as evidence of a robust predictive signal when accuracies remained close to zero.

### Heritability and variance structure in NIRS and metabolomic data

We estimated feature-wise heritability and variance components to separate genotypic from residual contributions to NIR and metabolomic features. Because location, year, block, and plot effects were not explicitly included in these feature-level models, the non-genetic component is reported as residual variation and is not attributed to environmental or genotype-by-environment effects. NIR wavelengths had low genotypic variance (mean 2%, maximum 17%) and low heritability (mean = 0.07, maximum = 0.46). Metabolomic features were more heritable (mean = 0.25), with several values above 0.90 (Figure S3). The same models provided feature-wise genotypic BLUPs, adjusted for sampling category, for subsequent prediction analyses.

### Model-specific prediction accuracy

Across the 24 traits, mean genomic predictive ability ranged from 0.375 for LASSO to 0.473 for rrBLUP (Table 1; Figure 1A-B), making rrBLUP the highest-mean model for genomic prediction. For phenomic prediction, mean predictive abilities were lower, ranging from 0.096 for rrBLUP to 0.215 for EN; EN was therefore the highest-mean phenomic model (Figure 1C-D). For metabolomic prediction, MBLUP yielded the highest mean predictive ability (0.424), followed by rrBLUP (0.408), whereas Random Forest yielded the lowest mean (0.350; Figure 1E-F). However, model rankings varied among traits and sampling sets, particularly for phenomic prediction. Sample-specific results for phenomic and metabolomic models are presented in Figures S4 and S5, respectively. Throughout the manuscript, the “highest-mean” model or sampling set refers to the one with the highest mean predictive ability within a predefined comparison. Such rankings are descriptive when predictive abilities are close to zero. In particular, phenomic model and sampling-set optima should not be interpreted as evidence of stable superiority or as breeding recommendations.

**Figure 1.**
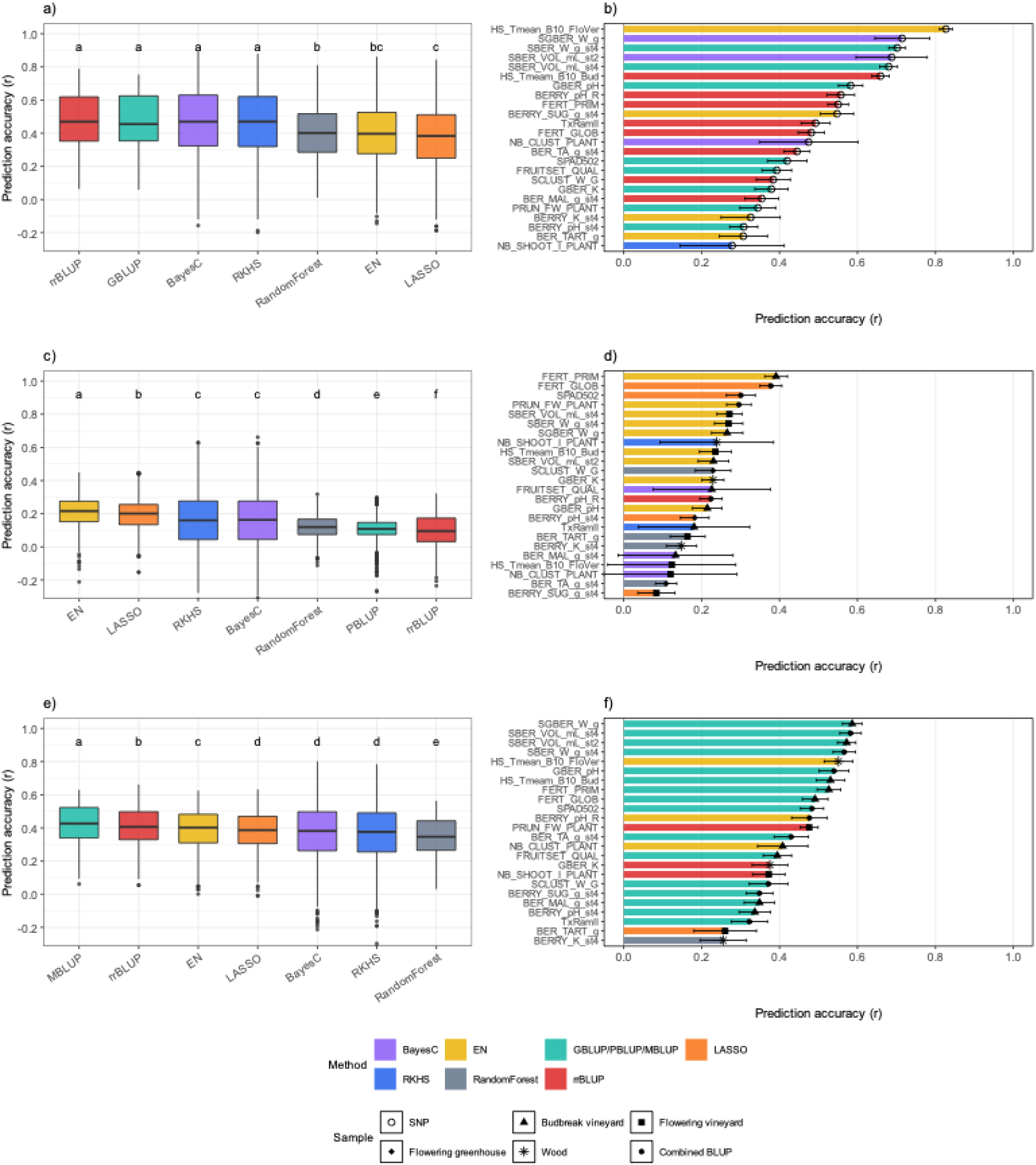
Performance of seven genomic, phenomic, and metabolomic prediction models in population 50025. Panel a) shows genomic predictive abilities across 24 traits, and panel b) the highest-mean genomic model for each trait. Panel c) shows phenomic predictive abilities after selecting the highest-mean sampling category for each trait and model, and panel d) the highest-mean phenomic model-sample combination for each trait. Panels e) and f) present the corresponding metabolomic results. Predictive ability is Pearson’s correlation coefficient (r) between observed and predicted phenotypic values. Letters above boxplots denote groups from a Kruskal-Wallis test followed by Dunn tests with Benjamini-Hochberg adjustment. Error bars in panels b), d), and f) are standard deviations across 50 prediction replicates. Colours denote methods: purple, BayesC; yellow, Elastic Net; turquoise, relationship-matrix BLUP; orange, LASSO; blue, RKHS; grey, Random Forest; and red, rrBLUP. The turquoise model is termed GBLUP, PBLUP, or MBLUP for genomic, phenomic, or metabolomic data, respectively. Black symbols denote predictor sources: open circles, SNPs; filled upward-pointing triangles, vineyard leaves at budbreak; filled squares, vineyard leaves at flowering; filled diamonds, greenhouse leaves at flowering; asterisks, wood; and filled circles, feature-wise BLUPs combining all sampling categories. BLUP, best linear unbiased prediction; LASSO, least absolute shrinkage and selection operator; RKHS, reproducing kernel Hilbert space; rrBLUP, ridge-regression BLUP.

**Table 1.** Prediction performance of models across omics layers in population 50025. Means and standard deviations are reported separately. For metabolomic and phenomic prediction, the sampling set yielding the highest mean predictive ability was selected independently for each trait and model. These selections should be interpreted descriptively when predictive abilities are close to zero. MBLUP and PBLUP denote the relationship-matrix BLUP models applied to metabolomic and phenomic data, respectively, distinguishing them from genomic GBLUP.

| Prediction models | Genomic prediction |  | Metabolomic prediction |  | Phenomic prediction |  |
| --- | --- | --- | --- | --- | --- | --- |
|  | Mean | Standard deviation | Mean | Standard deviation | Mean | Standard deviation |
| BayesC | 0.469 | 0.202 | 0.371 | 0.169 | 0.157 | 0.173 |
| EN | 0.396 | 0.183 | 0.392 | 0.117 | 0.215 | 0.093 |
| GBLUP/MBLUP/PBLUP | 0.470 | 0.158 | 0.424 | 0.118 | 0.109 | 0.072 |
| LASSO | 0.375 | 0.196 | 0.380 | 0.114 | 0.200 | 0.095 |
| RKHS | 0.466 | 0.197 | 0.367 | 0.172 | 0.157 | 0.172 |
| RandomForest | 0.402 | 0.156 | 0.350 | 0.107 | 0.120 | 0.070 |
| rrBLUP | 0.473 | 0.160 | 0.408 | 0.109 | 0.096 | 0.097 |

### Sampling effects on phenomic and metabolomic predictions

Four sampling categories were evaluated: wood, vineyard leaves at budbreak, vineyard leaves at flowering, and greenhouse leaves at flowering. We also calculated feature-wise BLUPs across the four categories. Each NIR spectrum contained 2,332 wavelengths. With the highest-mean model selected for each trait and sample, phenomic sample-level means were 0.172 for vineyard leaves at flowering, 0.161 for vineyard leaves at budbreak, 0.157 for combined-sample BLUPs, 0.145 for wood, and 0.088 for greenhouse leaves at flowering (Table 2; Figure S6A). These small differences were trait dependent and often accompanied by substantial variability, so the apparent sample ranking is descriptive. The highest trait-specific phenomic value was r = 0.391 for primary fertility (FERT_PRIM), obtained with EN and vineyard leaves at budbreak (Figure 1D; Figure S6B).

**Table 2.** Prediction performance by sampling category and omics layer in population 50025. Means and standard deviations are reported in separate columns. For each trait and sampling category, the prediction model with the highest mean was retained; phenomic rankings are descriptive when predictive abilities are close to zero.

| Sample | Metabolomic prediction | Phenomic prediction |
| --- | --- | --- |

|  | Mean | Standard deviation | Mean | Standard deviation |
| --- | --- | --- | --- | --- |
| Sample BLUP | 0.421 | 0.130 | 0.157 | 0.147 |
| Wood | 0.295 | 0.101 | 0.145 | 0.120 |
| Leaves at bud break | 0.382 | 0.133 | 0.161 | 0.125 |
| Leaves at flowering in the greenhouse | 0.199 | 0.145 | 0.088 | 0.121 |
| Leaves at flowering in the vineyard | 0.355 | 0.115 | 0.172 | 0.110 |

For metabolomic prediction, combined-sample BLUPs gave the highest sample-level mean (0.421), followed by vineyard leaves at budbreak (0.382), vineyard leaves at flowering (0.355), wood (0.295), and greenhouse leaves at flowering (0.199; Table 2; Figure S6C). The highest trait-specific metabolomic value was r = 0.586 for green-berry weight (SGBER_W_g), obtained with MBLUP and vineyard leaves at budbreak; berry weight and volume traits were among the most predictable across several sampling categories (Figure 1F; Figure S6D).

Parallel leaf sampling at flowering in the vineyard and greenhouse was used to assess whether controlled-environment profiles could predict vineyard phenotypes. For phenomic prediction, the relative performance depended on the trait. Greenhouse spectra exceeded vineyard spectra for berry sugar content (0.190 versus 0.054), cluster weight (0.298 versus 0.043), and, slightly, berry pH at stage 4 (0.126 versus 0.114). Vineyard spectra were higher for budbreak timing (0.334 versus -0.033), green-berry potassium (0.206 versus -0.042), and green-berry pH (0.112 versus 0.026; Figure 2A; Table S2). Given the low overall phenomic signal and large standard deviations for several traits, these trait-specific reversals are interpreted as descriptive rather than as evidence for a generally superior greenhouse sample.

**Figure 2.**
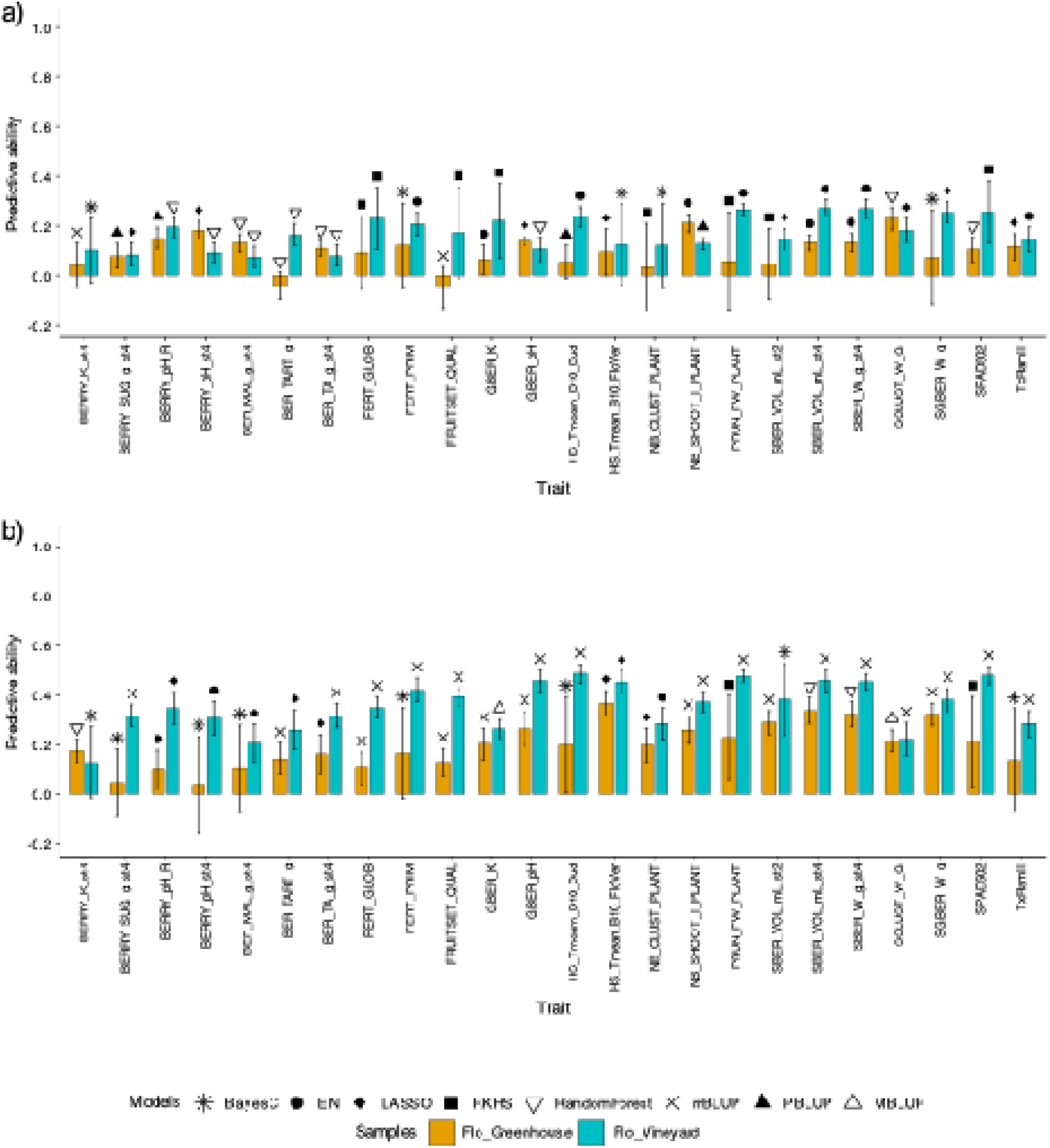
Effect of flowering-leaf sampling environment on phenomic and metabolomic prediction in population 50025. Panel a) compares trait-specific phenomic predictive abilities from leaves collected at flowering in the greenhouse and vineyard; panel b) presents the corresponding metabolomic predictions. For each trait and environment, the model with the highest mean was retained. Orange bars denote greenhouse samples and turquoise bars denote vineyard samples. Black symbols identify the selected model: asterisks, BayesC; filled circles, Elastic Net; filled diamonds, LASSO; filled squares, RKHS; open downward-pointing triangles, Random Forest; diagonal crosses, rrBLUP; filled upward-pointing triangles, PBLUP; and open upward-pointing triangles, MBLUP. Error bars are standard deviations across 50 prediction replicates. Predictive ability is Pearson’s correlation coefficient (r) between observed and predicted phenotypic values. MBLUP, metabolomic relationship-matrix best linear unbiased prediction; PBLUP, phenomic relationship-matrix best linear unbiased prediction.

For metabolomic prediction, vineyard leaves generally produced higher accuracies than greenhouse leaves. For example, berry sugar content increased from r = 0.041 in the greenhouse to 0.318 in the vineyard, and green-berry pH from 0.265 to 0.458. Berry potassium at stage 4 was an exception (0.175 in the greenhouse versus 0.1 in the vineyard). Greenhouse metabolomic accuracy reached r = 0.367 for flowering-to-veraison thermal time, showing that controlled-environment profiles retained predictive information for some traits (Figure 2B; Table S2).

### Comparison between genomic, phenomic, and metabolomic prediction models

To compare omics layers on the same individuals, GP, PP and MP were evaluated for the 150 individuals selected for metabolomic profiling, with trait-specific sample-model combinations chosen by highest mean. Genomic prediction was highest for most traits, metabolomic prediction was intermediate, and phenomic prediction was lowest (Figure 3; Table S3). The maximum genomic predictive ability was r = 0.827 for flowering-to-veraison thermal time (HS_Tmean_B10_FloVer) with EN. Metabolomic prediction reached r = 0.586 for green-berry weight (SGBER_W_g) with MBLUP and vineyard leaves at budbreak, whereas phenomic prediction reached r = 0.391 for primary fertility (FERT_PRIM) with EN and vineyard leaves at budbreak. Metabolomic prediction equaled or exceeded genomic prediction for a limited set of traits, including BERRY_pH_st4, FERT_GLOB, FRUITSET_QUAL, NB_SHOOT_I_PLANT, PRUN_FW_PLANT, and SPAD502. Because phenomic values were often low and variable, their trait-specific maxima are descriptive.

**Figure 3.**
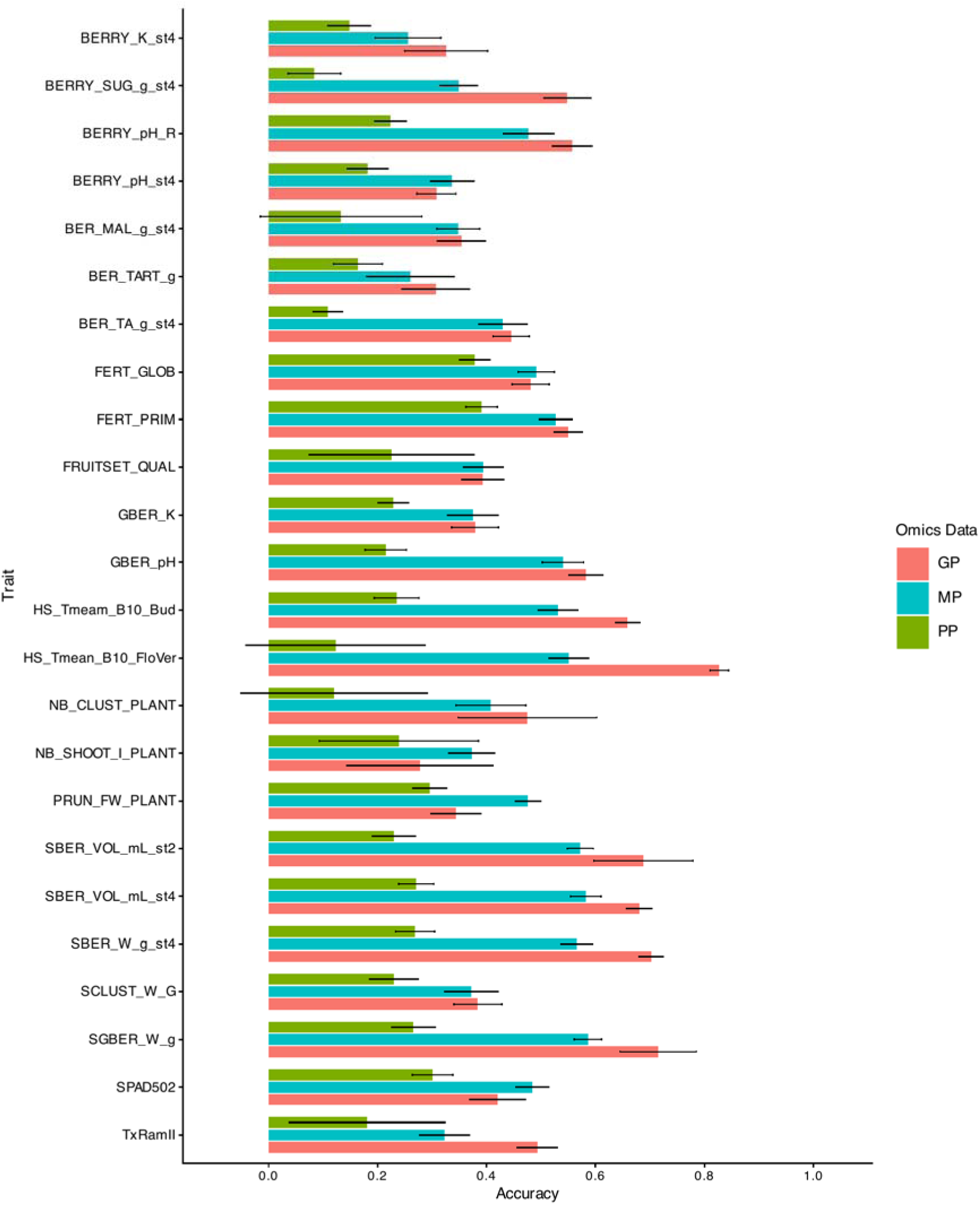
Predictive abilities of genomic, phenomic, and metabolomic models in population 50025. For each of the 24 traits and omics layers, the model-sample combination with the highest mean was retained. Coral, turquoise, and olive-green bars denote genomic, metabolomic, and phenomic prediction, respectively. Error bars are standard deviations across 50 prediction replicates. Predictive ability is Pearson’s correlation coefficient (r) between observed and predicted phenotypic values. Phenomic rankings should be interpreted descriptively when predictive abilities are close to zero. GP, genomic prediction; MP, metabolomic prediction; PP, phenomic prediction.

### Multi-omics prediction

Multi-omics models combined genomic (GP), phenomic (PP), and metabolomic (MP) inputs using BayesC or multi-kernel RKHS. GP and the genomic-inclusive combinations GP_MP, GP_PP, and GP_PP_MP formed the highest global statistical group, whereas MP, PP_MP, and PP formed successively lower groups (Kruskal-Wallis test followed by Dunn pairwise tests; Figure 4A). Thus, multi-omics integration did not provide a consistent global improvement over GP. The two-layer comparisons are detailed in Figure S7. GP_MP produced trait-specific gains in some cases, reaching 0.526 for FERT_GLOB versus 0.481 for GP and 0.491 for MP, and 0.411 for GBER_K versus 0.379 and 0.375, respectively (Figure S7B). In contrast, adding PP often reduced predictive ability for strongly genomic traits; for example, GP_PP reached 0.703 for HS_Tmean_B10_FloVer compared with 0.827 for GP (Figure S7A). PP_MP generally performed between MP and PP, indicating that adding phenomic information did not consistently improve predictions over MP alone (Figure S7C). Combining all three layers rarely improved on the highest-mean single- or two-layer model, although SPAD502 reached 0.509 with GP_PP_MP versus 0.484 with MP and 0.420 with GP (Figure 4B; Table S4). Overall, we found no evidence that multi-omics integration systematically improved predictive ability at the present sample size.

**Figure 4.**
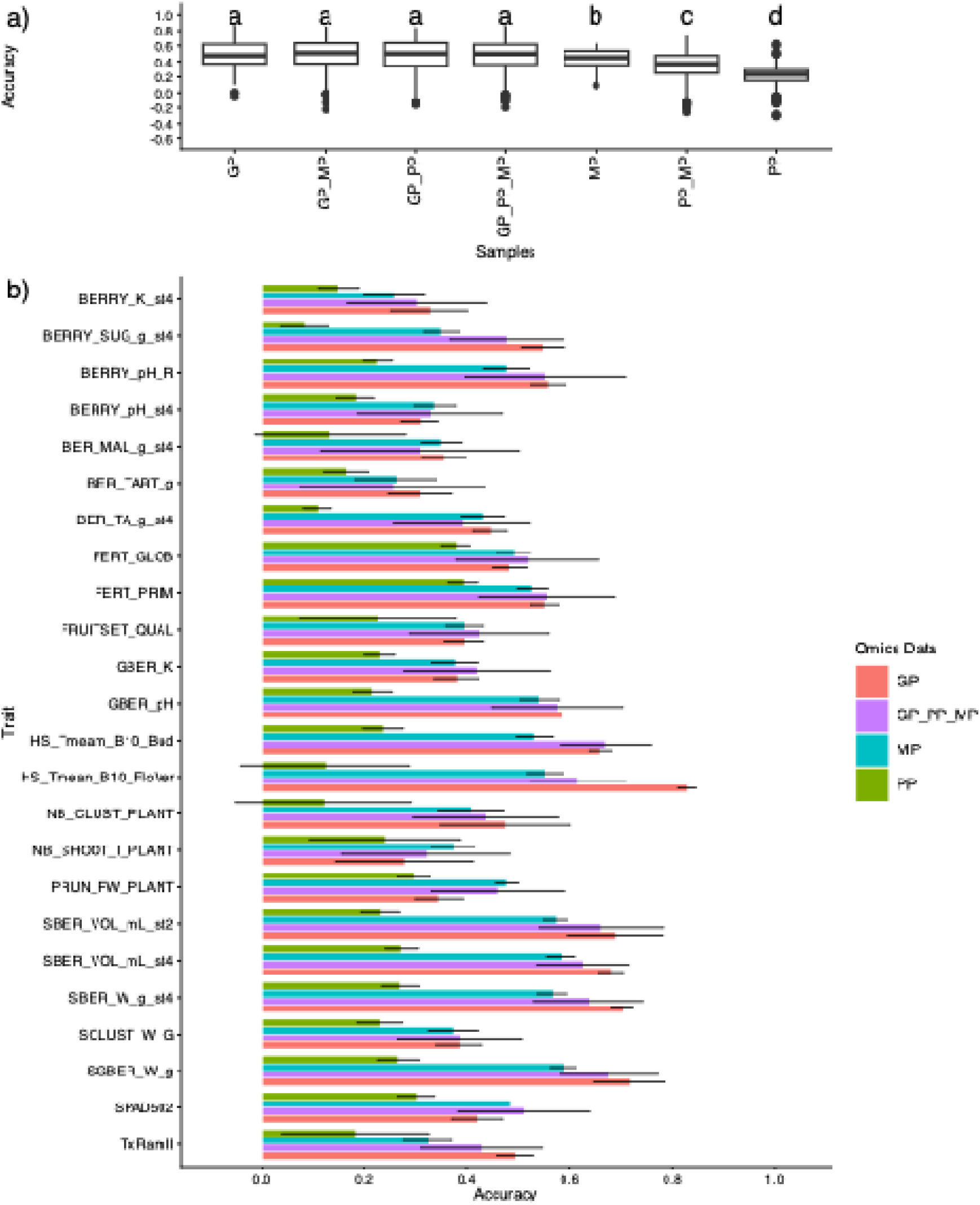
Performance of single- and multi-omics prediction models in population 50025. Panel a) shows predictive-ability distributions across 24 traits for single-layer and integrated models; panel b) shows trait-specific means for genomic, metabolomic, phenomic, and three-layer prediction. For each trait and data-layer configuration, the model-sample setting with the highest mean was retained. Letters above boxplots denote groups from a Kruskal-Wallis test followed by Dunn tests with Benjamini-Hochberg adjustment. Colors denote data layers: coral, genomic prediction (GP); turquoise, metabolomic prediction (MP); olive green, phenomic prediction (PP); purple, three-layer prediction (genomic+phenomic+metabolomic, GP_PP_MP). Error bars in panel b) are standard deviations across 50 prediction replicates. Predictive ability is Pearson’s correlation coefficient (r).

### Across-year transfer within population 50025

To assess temporal transfer, models were fitted using phenotypes recorded in one year (2019, 2020, or 2021) and evaluated against phenotypes recorded in another year for the same genotypes. For example, the 2019–2020 scenario denotes model fitting using 2019 phenotypes followed by evaluation using 2020 phenotypes. The SNP, NIR, and LC–MS predictor matrices and the genotype set remained unchanged across scenarios; only the phenotypic values used for model fitting and evaluation differed. Multi-year phenotypic BLUPs were also considered as either the training or evaluation phenotype. No genotypes were held out in this analysis. It therefore measures temporal correspondence within a fixed genotype set rather than predictive performance in new individuals or a formal estimate of genotype-by-year interaction. For each trait and omics layer, the highest-mean eligible model and, for phenomic and metabolomic prediction, sampling set were retained. Across the six year-to-year scenarios, mean predictive ability ranged from 0.418 to 0.518 for genomic prediction, from 0.438 to 0.528 for metabolomic prediction, and from 0.331 to 0.379 for phenomic prediction. Metabolomic prediction was slightly higher than genomic prediction when models were fitted using 2019 or 2020 phenotypes, whereas genomic prediction was slightly higher when models were fitted using 2021 phenotypes. Scenarios in which multi-year BLUPs were used for fitting or evaluation produced higher values across all three omics layers. However, because these BLUPs integrate observations from 2019-2021, including the year-specific records used in the comparisons, these results are not independent temporal validations and should be interpreted as measures of agreement with the multi-year phenotype (Figure S8; Table S5).

### Within- and across-population prediction

Within-population prediction was compared among 50025 (up to 250 individuals), 44910 (242), and RIxGW (254). Marker densities differed substantially: 7,386 SNPs in 50025, 18,867 in 44910, and 47,820 in RIxGW. Mean genomic predictive ability was highest in 50025 (0.520), followed by RIxGW (0.432) and 44910 (0.377; Table 3). Trait-specific exceptions included berry potassium and tartaric acid, for which 44910 was highest, and berry pH and pruning wood weight, for which RIxGW was highest (Figure 5A; Table S6). Phenomic prediction was available for 50025 and 44910 and had similar overall means (0.189 and 0.186, respectively), with population-specific differences among traits (Figure 5B; Table S6).

**Figure 5.**
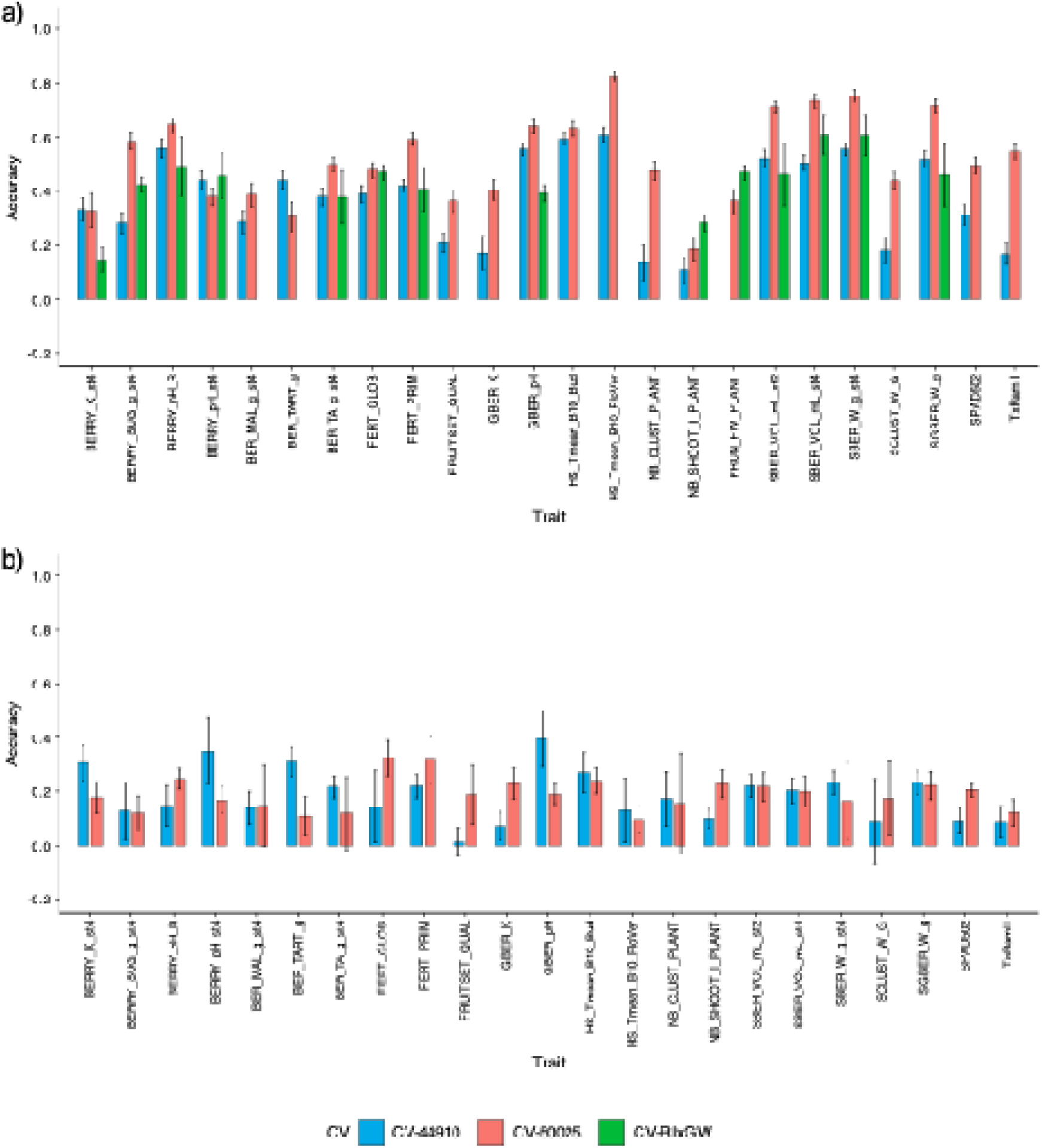
Within-population cross-validation across three biparental grapevine populations. Panel a) shows genomic prediction in populations 44910, 50025, and RIxGW; panel b) shows phenomic prediction in populations 44910 and 50025. For each trait and population, the eligible model with the highest mean was retained, together with the highest-mean sampling category for phenomic prediction. Blue, coral, and green bars denote populations 44910, 50025, and RIxGW, respectively. Error bars are standard deviations across 50 prediction replicates. Predictive ability is Pearson’s correlation coefficient (r) between observed and predicted phenotypic values. CV, cross-validation.

**Table 3.** Mean predictive ability and standard deviation for within- and across-population scenarios. Across-population labels use training-test notation (for example, 50025-44910 denotes training in 50025 and testing in 44910); CV denotes repeated within-population cross-validation.

| Prediction scenario | Genomic prediction |  | Phenomic prediction |  |
| --- | --- | --- | --- | --- |
|  | Mean | Standard deviation | Mean | Standard deviation |
| 44910-50025 | 0.181 | 0.086 | 0.089 | 0.042 |
| 44910-RIXGW | 0.048 | 0.057 | NA |  |
| 50025-44910 | 0.209 | 0.095 | 0.113 | 0.029 |
| 50025-RIXGW | 0.108 | 0.074 | NA |  |
| CV-44910 | 0.377 | 0.161 | 0.186 | 0.097 |
| CV-50025 | 0.520 | 0.165 | 0.189 | 0.060 |
| CV-RIXGW | 0.432 | 0.118 | NA |  |
| RIXGW-44910 | 0.090 | 0.060 | NA |  |
| RIXGW-50025 | 0.114 | 0.109 | NA |  |

Across-population prediction was consistently less accurate than within-population cross-validation (Table 3; Figure 6; Figure S9). Scenario labels use training-test notation: for example, 50025-44910 means training in 50025 and validation in 44910. Genomic mean predictive ability was 0.209 for 50025-44910 and 0.181 for the reciprocal 44910-50025 scenario, compared with within-population means of 0.520 and 0.377. Transfers involving RIxGW were lower (0.048-0.114). For phenomic prediction, 50025-44910 averaged 0.113 and 44910-50025 averaged 0.089, compared with within-population means of 0.189 and 0.186. Figure 6 focuses on scenarios in which 50025 was either the training or testing population, whereas Figure S9 groups all scenarios by testing population. These results show limited transferability across genetic backgrounds and should not be attributed specifically to environmental effects because population, relatedness, trial conditions, and marker sets are confounded.

**Figure 6.**
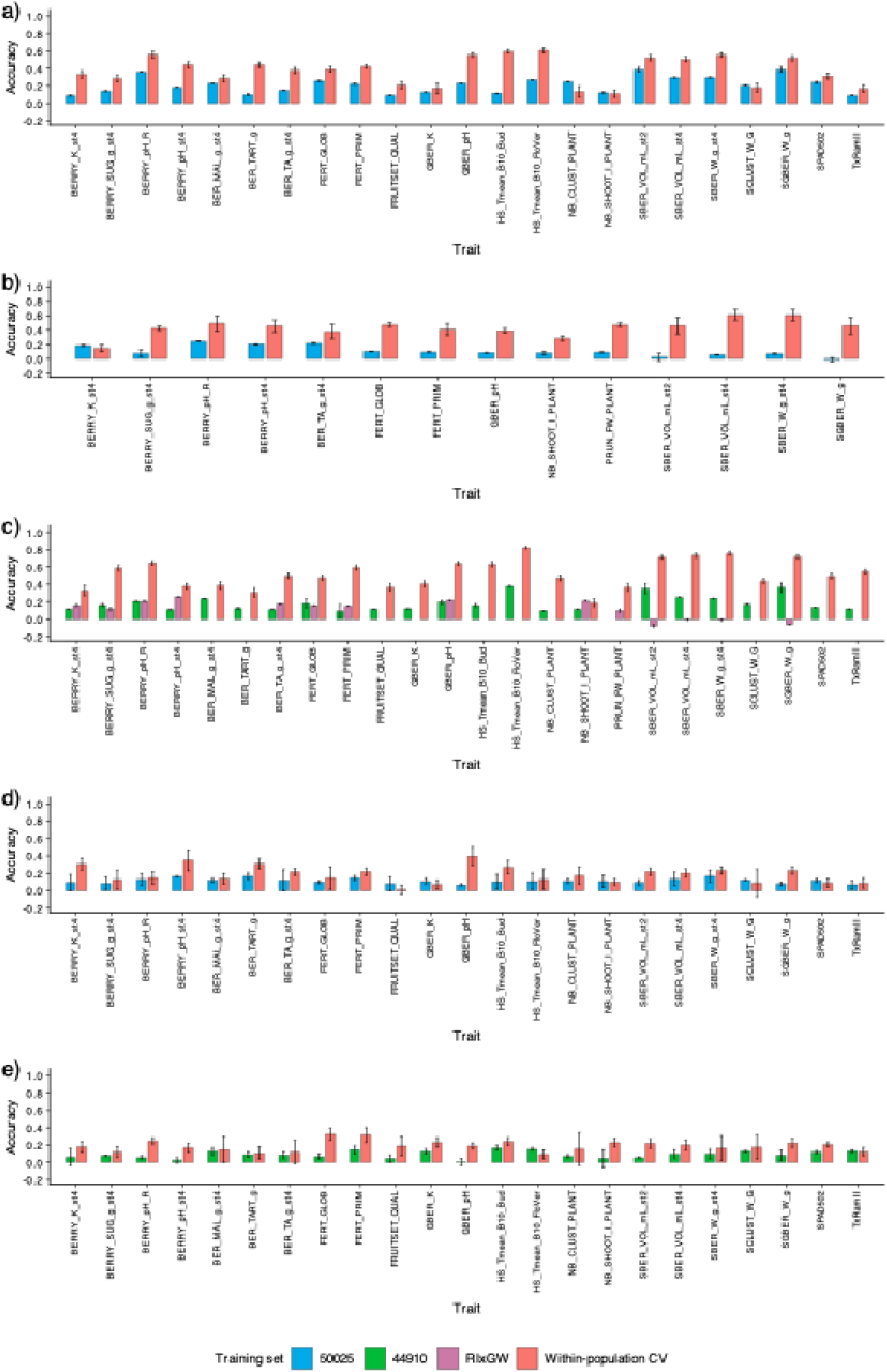
Genomic and phenomic prediction scenarios involving population 50025, grouped by testing population. Panel a) shows genomic prediction with population 44910 as the testing set; panel b) genomic prediction with RIxGW as the testing set; panel c) genomic prediction with population 50025 as the testing set; panel d) phenomic prediction with population 44910 as the testing set; and panel e) phenomic prediction with population 50025 as the testing set. Scenario direction is from the training population to the testing population. Blue, green, and magenta bars denote training with populations 50025, 44910, and RIxGW, respectively; coral bars denote within-population cross-validation for the relevant testing population. Error bars are standard deviations across 50 prediction runs when replication was performed; deterministic across-population fits may have a standard deviation of zero. Predictive ability is Pearson’s correlation coefficient (r). CV, cross-validation.

## Discussion

This study compared genomic, phenomic, and metabolomic prediction across models, sampling categories, and validation scenarios in grapevine breeding populations. Genomic prediction provided the highest and most stable signal for most traits. Metabolomic prediction was intermediate and competitive for selected traits, whereas phenomic prediction was generally low and variable. The results also showed that adding omics layers does not guarantee improvement and that cross-population transfer remains substantially less accurate than within-population validation.

Feature-wise variance partitioning showed that NIR wavelengths were dominated by residual variation, with mean heritability of 0.07, whereas metabolites were more heritable (mean = 0.25; maximum >0.90). Because location, year, block, and plot were not explicitly fitted, this residual component cannot be assigned to environmental or genotype-by-environment effects. The contrast nevertheless indicates stronger genetic determination of metabolomic features than of NIR wavelengths under our sampling design. Similar differences have been reported in other crops^23,24^. The low NIR heritability may reflect the heterogeneous tissues and phenological stages sampled; more homogeneous designs have yielded higher values in grapevine^19^ and maize^25^. Signals toward the mid-infrared can also be sensitive to temperature, moisture, and tissue scattering^26^. Accordingly, NIR spectra should be viewed here as integrative but weakly genetic predictors, whereas metabolomic profiles provided a more heritable molecular signature.

The Savitzky-Golay first derivative reduced baseline drift and improved prediction for some samples and models^27,28^. However, several phenomic means remained close to zero with large standard deviations, so the magnitude and direction of preprocessing effects should not be overinterpreted. Preprocessing remains an important technical choice, but it cannot compensate for weak or unstable predictive signal.

Model performance depended jointly on the omics layer and the target trait, although differences among methods were generally modest. For genomic prediction, rrBLUP, GBLUP, BayesC, and RKHS yielded similar mean predictive abilities (0.466-0.473), supporting the suitability of shrinkage, relationship-matrix, Bayesian, and kernel approaches for traits with predominantly polygenic architectures^14^. Nevertheless, no model class consistently dominated across traits. Penalized regression may be advantageous when phenotypic variation is influenced by a limited number of moderate- or large-effect loci. For example, EN yielded the highest predictive ability for HS_Tmean_B10_FloVer, a phenological trait associated with a major QTL on chromosome 16^29^. Traits related to berry acidity, potassium content, and pH have more complex genetic architectures^4,30^, but their model rankings did not consistently favor a single ridge-based or variable-selection approach. This agrees with comparisons in grapevine^19,31^ and other crops^32,33^ showing that relative model performance depends on genetic architecture, sample size, marker density, and population structure. The influence of the statistical model also differed among omics layers. MBLUP yielded the highest mean metabolomic predictive ability (0.424), whereas EN had the highest phenomic mean (0.215). Metabolomic prediction generally occupied an intermediate position between genomic and phenomic prediction, although model rankings varied among traits and sampling sets. For phenomic prediction, predictive abilities were low and variable, and no consistent relationship emerged between model ranking and trait genetic architecture, as also observed by Robert et al.^34^. Consequently, the highest-mean phenomic models should be interpreted descriptively rather than as evidence of stable model superiority.

Sampling stage and tissue should be considered biological filters that determine which components of genotype-dependent variation are captured. LC-MS profiles provide a relatively direct but dynamic representation of metabolism and are therefore sensitive to developmental stage and environmental conditions. In grapevine, phenology is a major source of variation in the leaf metabolome, together with temporal and spatial variation during sampling^35^. The greater informativeness of vineyard leaves collected at budbreak relative to the other individual sampling sets may therefore reflect genotype-dependent differences in spring metabolic reactivation and source-sink transitions. Early shoot and inflorescence development relies substantially on the mobilization of stored carbon and nitrogen until approximately flowering^36^, potentially connecting early-season leaf chemistry with subsequent variation in vegetative growth, fertility, and berry development. This interpretation remains a hypothesis because developmental stage and sampling environment were not experimentally separated. The performance of the combined-sample BLUPs nevertheless suggests that extracting the component shared across sampling stages can reduce stage-specific variation while retaining metabolomic differences associated with genotype. The biological information captured by NIR spectra differs from that measured by LC-MS. A NIR spectrum represents an integrated chemical and physical fingerprint of a tissue rather than the abundance of specific metabolites; consequently, a more metabolically active tissue is not necessarily a better substrate for phenomic prediction. In wheat, poplar, and grapevine, the relative value of leaf, grain, and wood spectra depended on the trait and experimental context^16,19^. Brault et al.^19^ found only modest differences between grapevine leaf and wood spectra, whereas combining tissues or years generally improved phenomic prediction. This is consistent with the absence of a stable phenomic advantage for any single tissue in the present study. Grapevine canes undergo marked reprogramming of phenylpropanoid metabolism around budbreak^37^. Wood spectra may therefore emphasize relatively persistent structural and storage-related variation, whereas leaf spectra capture a more dynamic physiological state. Combining spectra across sampling stages may help retain genotype-persistent information from both components, but the low and variable phenomic predictive abilities observed here require this hypothesis to be tested in independent populations and environments.

The contrast between vineyard and greenhouse sampling further indicates that the value of proxy-omic information depends on the biological correspondence between the sampling and target environments. The lower overall performance of greenhouse profiles suggests that controlled conditions did not reproduce all physiological variation relevant to field phenotypes. Nevertheless, their predictive value for selected traits indicates that some genotype-associated biochemical variation persists across environments. Metabolic profiles collected from young maize roots under controlled conditions have similarly predicted hybrid performance in field trials^38^, demonstrating that early controlled-environment measurements can be informative when they capture processes shared with the target phenotype. Greenhouse sampling could therefore support early screening in grapevine, but only for trait-sample associations that are reproducible across populations and growing cycles. Aggregating data across tissues and years through BLUP estimation increased predictive accuracy, likely by isolating the stable genetic component from environmental noise, as also observed in previous studies in grapevine or in rice and maize^19,39,40^.

While numerous studies have compared genomic and phenomic or genomic and metabolomic predictions this work is, to our knowledge, the first to simultaneously benchmark genomic, phenomic, and metabolomic prediction in grapevine. The relative performance of the three omics layers reflects differences in both their proximity to the phenotype and their temporal stability. SNPs provide stable inherited information, whereas metabolite and NIR profiles represent downstream molecular and biochemical states influenced by genotype, development, and environment. Metabolites can therefore act as intermediate phenotypes that integrate multiple regulatory processes more directly related to complex traits than individual markers. This may explain why metabolomic prediction approached or exceeded genomic prediction for selected traits, consistent with studies of fruit quality in tomato and hybrid performance in maize^41,42^. However, such agreement does not demonstrate that metabolomic profiles specifically capture genotype-by-environment interactions; establishing that contribution would require replicated multi-environment sampling and models that explicitly estimate environmental and interaction effects.

NIR-based phenomic prediction occupies a different operational niche. Its main advantage is the rapid and inexpensive acquisition of high-dimensional biochemical fingerprints rather than consistently high predictive ability^16,19^. The variable performance reported across tissues, traits, populations, and environments in grapevine and other crops indicates that spectral similarity is not a universal surrogate for genomic relatedness^43–46^. Phenomic prediction is therefore most relevant when a robust positive signal has been demonstrated for a defined tissue, developmental stage, and target population. Rankings derived from near-zero predictive abilities should not be used to identify a preferred sampling strategy or to support breeding decisions.

The absence of a systematic gain from multi-omics integration is biologically informative. Adding a data layer can improve prediction only when it contributes information that is both complementary to the existing layers and estimable at the available sample size. Metabolomic profiles may partly mediate genetic effects and thus overlap with information already captured by SNPs, whereas a weak phenomic signal can increase estimation variance without adding useful information. Previous studies have likewise reported that multi-omics integration is beneficial for some traits but neutral or detrimental for others^19,32,40,42,47^. With only 150 individuals available for metabolomic and integrated analyses, the simultaneous estimation of multiple high-dimensional effects was challenging. These results favor trait-specific integration guided by independent validation rather than the general assumption that adding omics layers will improve prediction.

The across-year analysis provides evidence of temporal correspondence but should not be interpreted as an independent assessment of genotype-by-year interaction. Models were fitted and evaluated using different year-specific phenotypes for the same genotype set, without holding out individuals. Prediction accuracy varied markedly among years, reflecting the hot and dry 2019, the cooler and stormy 2020, and the humid 2021 vintages. Multi-year phenotypic BLUPs provided higher accuracies than single-year datasets, confirming that multi-year training improves predictive reliability. However, those BLUPs incorporate information from the annual records against which they were compared, accounting in part for their stronger associations. Explicit assessment of temporal stability would require independent genotype validation within multiple environments and models that partition main genetic, environmental, and interaction effects^43,48^. Such designs would also clarify whether metabolomic or spectral profiles improve prediction by capturing stable genetic differences, environment-specific responses, or both.

The decline in predictive ability across populations is consistent with the dependence of genomic prediction on relatedness, allele frequencies, and linkage-phase persistence^11,49,50^. The comparatively greater transfer between the two interspecific populations may reflect their closer genetic relationship, whereas RIxGW represents a more distinct *V. vinifera* background. However, population, trial environment, phenotyping year, rootstock, and marker density varied simultaneously, preventing their individual contributions from being resolved. Expanding the training set will therefore not necessarily improve transfer unless the additional individuals represent the allelic diversity and haplotypes present in the target population. Optimized multi-population training sets that balance diversity with relatedness offer a more defensible strategy^9,43^.

For operational breeding, genomic selection consequently remains the most broadly reliable approach evaluated here. Metabolomic prediction may be valuable for traits for which it provides reproducible information beyond genomic prediction and for which the additional assay cost is justified. Phenomic prediction offers a lower-cost and higher-throughput option, but its use should be restricted to validated trait and sampling combinations. A staged strategy in which an inexpensive phenomic screen reduces the number of candidates subsequently evaluated by genomic or metabolomic methods is conceptually attractive^16,19,43^. Its practical value, however, must be established prospectively by comparing selection response, retained genetic diversity, cost per selected genotype, and breeding- cycle duration with those of genomic selection alone.

This study provides an integrated comparison of genomic, phenomic, and metabolomic prediction in grapevine breeding populations. Genomic prediction was most accurate for most traits, metabolomic prediction was intermediate and occasionally competitive, and phenomic prediction, despite its low marginal cost and high throughput, produced the weakest and least stable signal. SNP genotyping requires a greater initial investment than NIR spectroscopy, but genomic data are stable and can be reused across traits, years, and environments. By contrast, LC-MS profiling entails substantial sample-processing and analytical costs and is therefore most relevant when it provides reproducible gains for high-value traits. Sampling category, statistical model, and relatedness between training and testing populations affected performance, whereas multi-omics integration produced no consistent global gain at the present sample size. These findings support genomic selection as the primary predictive framework, complemented by trait-specific metabolomic or phenomic screening when the additional predictive value, throughput, and cost per selected genotype justify its use.

## Methods

### Plant material and experimental designs

The focal population (code 50025) derives from a cross between IJ119 (female; INRAE-ResDur program) and Divona (male; Agroscope, Switzerland), both carrying resistance to downy and powdery mildews. The cross yields 250 progenies with complex interspecific ancestry (North American, Asian, and European *Vitis*), diverse resistance-gene combinations, and breeding-relevant segregation. A full description of the population, experimental layouts, phenotyping, and genotyping is provided in Chedid (2023)^21^. In brief, two complementary designs were established in the vineyard at immediate proximity of the INRAE center in Colmar, France: (i) a greenhouse collection (n = 250) maintained primarily for germplasm preservation; and (ii) a vineyard trial (n = 209) planted as three plants per genotype, replicated in two blocks, one unsprayed to assess disease resistance, and one protected to measure agronomic traits. Only the protected block has been used in this manuscript.

To assess across-population prediction, two further breeding populations were used:

- 44910 (n = 242): a cross HYB.42103.Col.2087N x Chardonnay. HYB.42103.Col.2087N is an interspecific hybrid (IJ119 x Bronner), originating from a regional program targeting DRVs with typicity aligned to Bourgogne/Champagne. This population was phenotyped in greenhouse conditions in 2020 and 2021 (agronomic traits).
- RIxGW (n = 254): a pure *V. vinifera* cross Riesling x Gewurztraminer, phenotyped in vineyard conditions at Bergheim, France in an Alsace appellation vineyard (agronomic traits).

All three populations were genotyped by a common pipeline (see below).

### Phenotyping

For population 50025, 24 agronomic traits covering phenology, vegetative growth, yield components, and berry composition were recorded from 2019 to 2021. Trait definitions and phenotyping years are provided in Table S7. Previously estimated multi-year phenotypic BLUPs were used where indicated^21,51,52^. Population 44910 was phenotyped for 23 agronomic traits in greenhouse conditions in 2020 and 2021, and RIxGW for 14 traits in vineyard conditions in 2017 and 2018.

### Genotyping and marker processing

Leaf DNA was genotyped by genotyping-by-sequencing using ApeKI digestion^53^. Reads were aligned to PN40024.v4^54^, and SNP discovery and filtering followed Stacks^55^. Quality thresholds were read depth DP ≥ 4 and allelic depth AD > 2. A MAF threshold of 0.005 was used to remove monomorphic or nearly monomorphic loci in these biparental populations, and missing genotypes were imputed with Beagle 5.5^56^. The within-population matrices contained 7,386 SNPs for 50025, 18,867 for 44910, and 47,820 for RIxGW. Across-population analyses used 3,831 SNPs shared by 50025 and 44910, 1,834 shared by 50025 and RIxGW, and 5,468 shared by 44910 and RIxGW.

### Sampling for NIR spectroscopy and metabolomics

To identify cost-effective sampling strategies for prediction, tissues were collected in 2023 from winter wood and leaves at defined phenophases: (i) wood at end of winter; (ii) vineyard leaves at end of budbreak; and (iii) leaves at end of flowering in both greenhouse and vineyard. For wood, one secondary branch per plot (per genotype) was sampled. For vineyard leaves, one leaf per plant (three per genotype) was collected; in the greenhouse, one leaf per branch (one to two per genotype). Greenhouse leaves at end of budbreak were also sampled for 44910. See Figure S10 for a schematic of sampling across tissues and time points.

### NIR spectroscopy: sample preparation, acquisition, and preprocessing

Wood samples were kept cold, then oven-dried three days at 70 °C and milled (electric pencil sharpener on internodes). Leaf samples were oven-dried three days at 70 °C; leaves per genotype were pooled and milled. Spectra were recorded with a Bruker’s TANGO FT-NIR spectrometer over the near-infrared region (approximately 800-2500 nm, i.e., 12,500-4,000 cm ¹). Each sampling produced a high-dimensional spectrum used as input features (wavelengths). To reduce baseline and slope effects and enhance informative absorbance features, spectra were transformed using the Savitzky-Golay first derivative^28^, consistent with prior work in grapevine phenomic selection^19^.

### Untargeted UHPLC-HRMS metabolomics: sample preparation, acquisition, and processing

Untargeted metabolomics was performed on 150 individuals from population 50025 selected using the CDmean criterion to maximize reference-set representativeness^57^. Leaf discs collected from the leaves used for NIR acquisition were stored at -80 °C, freeze-dried, and ground with a TissueLyser II bead mill (Qiagen, Courtaboeuf, France). Powder (10 mg) was extracted with methanol (50 μL/mg) containing chloramphenicol as an internal standard (5 mg/L)^58^, sonicated for 10 min, and centrifuged at 12,000 x *g* and 10 °C for 10 min. Wood samples were processed similarly but underwent microwave extraction before sonication and two successive extractions with equal methanol volumes (35 μL/mg per extraction). After each extraction, samples were sonicated and centrifuged, and the supernatants were pooled. Extracts (1 μL) were analyzed using a Vanquish Flex binary UHPLC system coupled to an Exploris 120 Q-Orbitrap HRMS (Thermo Scientific). Separation was performed at 30 °C on an Acquity HSS T3 column (100 x 2.1 mm, 1.8 μm, 100 Å; Waters) at 0.33 mL/min. Eluents were acetonitrile/0.1% formic acid (A) and water/0.1% formic acid (B). The gradient was 85% B at 0-1 min, 85-70% B at 1-4 min, 70-50% B at 4-5 min, 50-40% B at 5-6.5 min, 40-1% B at 6.5-8 min, 1% B at 8-10 min, and 1-85% B at 10-11 min. The HRMS used heated electrospray ionization in positive and negative modes with the following settings: spray voltage, ±3.0 kV; sheath, auxiliary, and sweep gas flows, 40, 10, and 1 arbitrary units, respectively; capillary and auxiliary-gas heater temperatures, 320 °C; full-MS resolution, 60,000 FWHM at *m/z* 200; and scan range, *m/z* 85–1,200. The EASY-IC source provided internal calibration across the full mass range. Data were processed in Compound Discoverer 3.3.3.200 (Thermo Scientific). Retention times were aligned using ChromAlign^59^; components were retained when peak rating was ≥4 in at least four samples; ions were searched across the dataset with a 5-ppm tolerance and a signal-to-background threshold of 1.5 for gap filling; and batch effects were corrected from quality-control samples using systematic error removal with random forest (SERRF; 200 trees).

### Feature-wise BLUPs and heritability (NIR wavelengths and metabolites)

For each feature (wavelength or metabolite) in population 50025, we fitted the following mixed model:

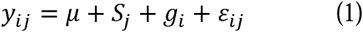

where y_ij_ is the preprocessed NIR value at one wavelength or the intensity of one LC-MS feature for genotype i in sampling category j, µ is the intercept, S_j_ is the fixed effect of sampling category, g_i_ is the random genotypic effect with g_i_ distributed as *N*(0, σ^2^_g_), and ε_ij_ is the residual distributed as *N*(0, σ^2^_e_). The four sampling categories were wood, vineyard leaves at budbreak, vineyard leaves at flowering, and greenhouse leaves at flowering. These categories represent distinct tissue, developmental-stage, and growing-environment combinations and were not biological replicates within a tissue x time-point combination. Genotypic BLUPs were extracted separately for each feature and used as combined-sample predictors.

Feature-wise reliability of the adjusted genotypic mean across sampling categories, expressed as an entry-mean broad-sense heritability (*H^2^*), was estimated as:

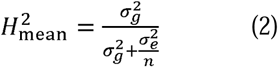

where σ^2^ and σ^2^ are the genotypic and residual variance components and n is the number of available sampling categories per genotype (up to four). The framework was applied separately to the NIR wavelength and LC-MS feature matrices.

### Prediction models

Seven single-omics frameworks were compared: rrBLUP, relationship-matrix BLUP (GBLUP for genomic, PBLUP for phenomic, and MBLUP for metabolomic data), BayesC, LASSO, Elastic Net, Random Forest, and RKHS. BayesC and RKHS were also extended to two- and three-layer multi-omics models. Analyses were conducted in R with glmnet^60^, rrBLUP^61^, BGLR^62^, and ranger^63^.

Linear ridge models (rrBLUP and relationship-matrix BLUPs): for rrBLUP, all standardized features were fitted simultaneously as random effects:

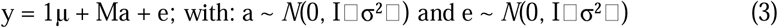

where y is the phenotype vector containing *n* observations, μ is the intercept, M is the *n × p* centered and scaled predictor matrix containing SNPs, preprocessed NIR wavelengths, or LC-MS features, a is the vector of *p* random feature effects, and e is the residual vector. I and I are identity matrices of dimensions *p × p* and *n × n*, respectively. The feature-effect variance (σ²) and residual variance (σ²) were estimated by restricted maximum likelihood (REML).

The relationship-matrix BLUP model was:

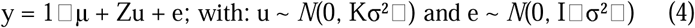

where Z links observations to genotypes, u is the vector of genotype effects, and K is the relationship or similarity matrix derived from the corresponding omics data. For the single-omics analyses in population 50025, each column of the predictor matrix M was centered and scaled, and the matrix was calculated as: K = MM / p, where p is the number of SNPs, NIR wavelengths, or LC-MS features. For the within- and across-population analyses, relationship matrices were calculated using the A.mat function of the R package rrBLUP. The relationship-matrix model is termed GBLUP for SNPs, PBLUP for NIR spectra, and MBLUP for metabolomic profiles. Variance components were estimated by REML.

Bayesian models (BayesC): BayesC assigns each standardized feature effect a two-component mixture prior comprising a point mass at zero and a Gaussian distribution for non-zero effects^8^. The model was:

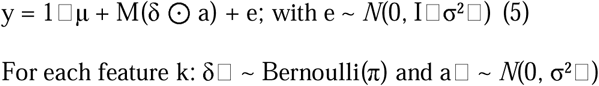

For each feature k: δ ∼ Bernoulli(π) and a ∼ *N*(0, σ²) where y is the phenotype vector, μ is the fixed intercept, M is the n × p matrix of centered and scaled SNPs, NIR wavelengths, or LC-MS features, a is the effect of feature k, and δ indicates whether that effect is included in the model. The effective feature effect is therefore δ a : it equals zero when δ = 0 and follows a Gaussian distribution when δ = 1. The parameter π, representing the proportion of non-zero effects, was assigned the default beta prior implemented in BGLR and estimated from the data.

Kernel model (RKHS implementation): the Reproducing Kernel Hilbert Space (RKHS) regression^64^ captures relationships between omics and phenotypes. For a single omics layer, the model was:

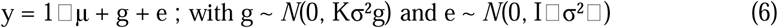

where y is the phenotype vector, μ is the intercept and the only fixed effect, g is the random kernel effect, and e is the residual vector. Each column of the n × p omics matrix M was centered and scaled, and the kernel matrix was calculated as: K = MM / p, where p is the number of SNPs, NIR wavelengths, or LC-MS features.

BayesC and RKHS models were fitted using the BGLR package with 20,000 MCMC iterations, of which the first 4,000 were discarded as burn-in; every fifth subsequent iteration was retained. Predictions were calculated from the posterior means based on the retained samples. Identical MCMC settings were used across omics layers and their combinations.

Penalized regression (LASSO and Elastic Net): LASSO^65^ and Elastic Net were implemented with glmnet. For both methods, the model was:

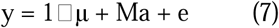

where y is the phenotype vector, μ is the intercept, M is the n × p matrix of predictors, a is the vector of feature effects, and e is the residual vector. Predictor columns were centered and scaled, whereas the intercept was not penalized. The coefficients were estimated by minimizing:

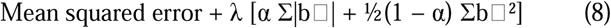

where λ controls the overall strength of regularization and α determines the balance between the L1 and L2 penalties. LASSO was fitted with α = 1, resulting in an exclusively L1 penalty that can set feature effects exactly to zero. EN was fitted with α = 0.5, giving equal weighting to the L1 and L2 components and allowing both feature selection and shrinkage of correlated predictors. For each trait and omics matrix, λ was selected by five-fold cross-validation using cv.glmnet. The value yielding the minimum mean cross-validation error, lambda.min, was retained for prediction.

Random Forest: Random Forest^66^ regression was fitted using the ranger package. Each forest comprised 100 regression trees. For each tree, individuals were sampled with replacement from the corresponding training set. At each node, mtry = floor(sqrt(p)) predictors were randomly selected as candidates for splitting, where p is the number of SNPs, NIR wavelengths, or LC-MS features. Splits were selected using the default variance-reduction criterion for regression, and the default minimum node size of five observations was used. For an individual i, the prediction was the mean of the predictions from the 100 trees:

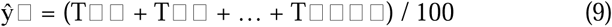

where T is the prediction from tree b. Forests were fitted exclusively using the individuals in each training fold and were then applied to the corresponding held-out individuals. No hyperparameter tuning or variable-importance calculation was performed.

Integrative models combined genomic (GP), phenomic (PP), and metabolomic (MP) layers in two- or three-layer configurations. Individual identifiers were aligned across the phenotype vector and all omics matrices before fitting, and each analysis was restricted to individuals represented in every layer included in that configuration. Each layer contributed an additive model component; cross-layer interaction terms were not fitted.

Multi-omics BayesC: for Q omics layers, the model was:

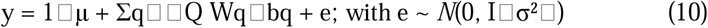

where μ is the intercept and the only fixed effect, Wq is the centred and scaled n × pq feature matrix for omics layer q, and bq is its vector of BayesC feature effects. For feature k in layer q, the prior was: bq,k = 0, with probability 1 – πq; bq,k ∼ *N*(0, σ²b,q), with probability πq. Each omics layer was specified as a separate BayesC component in BGLR and therefore had its own inclusion probability πq and non-zero effect variance σ²b,q. The inclusion probabilities were assigned the default beta priors implemented in BGLR and estimated from the data.

Multi-kernel RKHS: for Q omics layers, the model was:

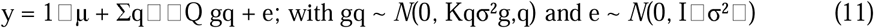

where gq is the random kernel effect of omics layer q, with a separate variance component σ²g,q. For each layer, the normalized linear similarity matrix was calculated as: Kq = WqWq / pq, where pq is the number of features in layer q. Thus, each omics layer contributed a separate linear kernel. No Gaussian kernel, nonlinear transformation, or bandwidth parameter was used.

Multi-omics BayesC and RKHS models were fitted using 20,000 MCMC iterations, with the first 4,000 discarded as burn-in and every fifth subsequent iteration retained. Predictions were calculated from the posterior means of the retained samples.

### Prediction scenarios and accuracy assessment

Three validation schemes were used:

- Repeated cross-validation: individuals were randomly partitioned using an 80:20 training-validation split corresponding to one held-out fifth, and the procedure was repeated 50 times. Prediction ability was Pearson’s r between observed and predicted phenotypes. Means and standard deviations were calculated across the 50 prediction replicates;
- Across-population prediction: models were trained in one population and applied to another. Labels use training-test notation (for example, 50025-44910 means training in 50025 and testing in 44910). Stochastic models were refitted 50 times; deterministic models can therefore have a standard deviation of zero.
- Across-year prediction: models were trained on one year-specific phenotype (2019, 2020, or 2021) or a multi-year phenotypic BLUP from 50025 and tested on another year-specific phenotype or BLUP for the same genotypes. No cross-validation between genotypes was used in this scenario, which evaluates temporal transfer for a fixed genotype set.

## Data availability

The raw datasets generated for this study are accessible on the data.gouv public repository at: https://doi.org/10.57745/NDN2LL.

## Additional information

Supplementary data to this article can be found online.

## Acknowledgements

This work was supported by funding through the IB_2023_MetabOptimum project of the INRAE BAP division, as well as the DIGIT-BIO travel grant, the OIV and the INRAE BAP division research grants for the PhD thesis of Clémentine Borrelli. We thank all the staff of the INRAE UEAV (Unité d’Expérimentation Agronomique et Viticole, doi: 10.15454/1.5483269027345498E12) experimental unit of Colmar for their support and the management of the vineyards. We thank Amandine Velt for her assistance in the use of the unit bioinformatics server.

## Author contribution

Borrelli C: Formal analysis, Investigation, Visualization, Writing - Original Draft. Delannoy L, Chepca H, Calcaterra M, and Chedid E: Formal analysis, Investigation. Arnold G and Dumas V: Resources. Baltenweck R, Maia-Grondard A, and Hugueney P: Methodology, Resources, Investigation, Formal analysis, Writing - Review & Editing. Merdinoglu D and Duchêne É: Conceptualization, Writing - Review & Editing. Avia K: Conceptualization, Investigation, Writing - Review & Editing, Supervision, Funding acquisition.

