## Supplementary data for "Comparative assessment of genomic, phenomic, and metabolomic prediction models in biparental grapevine breeding populations"

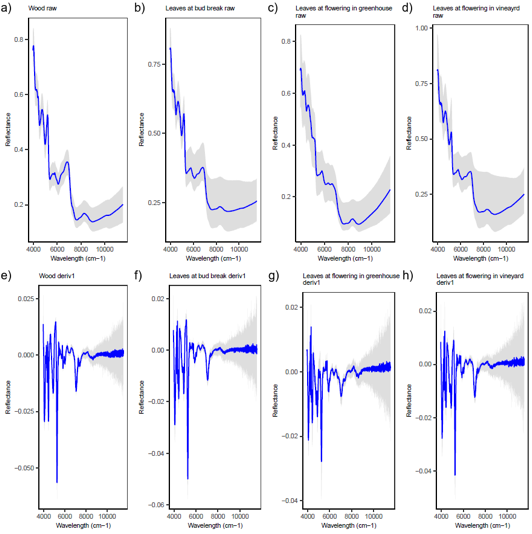

**Figure S1.** Effect of first-derivative preprocessing on near-infrared spectra from population 50025. Panels a), b), c), and d) show raw spectra from wood, vineyard leaves at budbreak, greenhouse leaves at flowering, and vineyard leaves at flowering, respectively. Panels e), f), g), and h) show the corresponding spectra after Savitzky-Golay first-derivative transformation. Blue lines represent mean spectra across individuals, and grey bands show the observed minimum-to-maximum range. NIR, near-infrared.

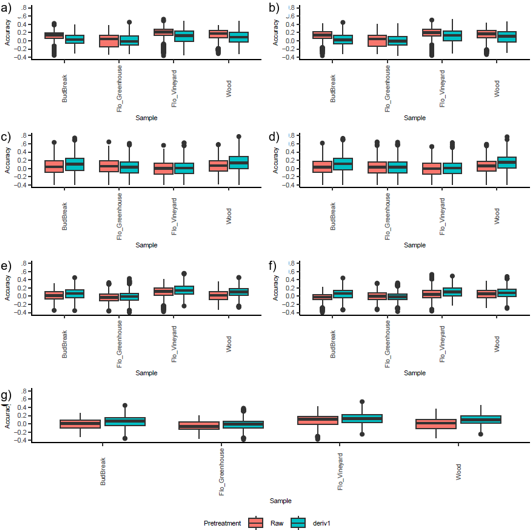

**Figure S2.** Effects of spectral preprocessing and sampling category on phenomic prediction in population 50025. Panels a)-g) show results for LASSO, Elastic Net, BayesC, RKHS, rrBLUP, Random Forest, and PBLUP, respectively. Coral and turquoise boxplots denote raw spectra and spectra transformed using the Savitzky-Golay first derivative, respectively. Sampling categories are vineyard leaves at budbreak, greenhouse leaves at flowering, vineyard leaves at flowering, and wood. Predictive ability is Pearson’s correlation coefficient (r). LASSO, least absolute shrinkage and selection operator; PBLUP, phenomic relationship-matrix best linear unbiased prediction; RKHS, reproducing kernel Hilbert space; rrBLUP, ridge-regression best linear unbiased prediction.

Table S1. Summary of pre-treatment effects on phenomic prediction accuracy for the 50025 population. Mean and standard deviation of prediction accuracies for each combination of sample, prediction model, and NIR spectra pre-treatment. Raw: untreated spectra, deriv1: spectra processed with the first Savitzky-Golay derivative.

| Pre-treatment of NIR spectra | Prediction models | Leaves at Bud Break | | Leaves at the Flowering in the Greenhouse | | Leaves at the flowering in the Vineyard | | Wood | |
| --- | --- | --- | --- | --- | --- | --- | --- | --- | --- |
|  |  | Mean | Standard deviation | Mean | Standard deviation | Mean | Standard deviation | Mean | Standard deviation |
| Raw | BayesC | 0.039 | 0.196 | 0.05 | 0.198 | -0.005 | 0.19 | 0.071 | 0.18 |
| deriv1 | BayesC | 0.109 | 0.208 | 0.02 | 0.201 | -0.001 | 0.183 | 0.135 | 0.217 |
| Raw | EN | 0.122 | 0.141 | 0.019 | 0.157 | 0.18 | 0.156 | 0.136 | 0.158 |
| deriv1 | EN | 0.037 | 0.141 | 0.011 | 0.147 | 0.121 | 0.152 | 0.099 | 0.164 |
| Raw | GBLUP | -0.005 | 0.116 | -0.034 | 0.115 | 0.085 | 0.157 | 0.008 | 0.141 |
| deriv1 | GBLUP | 0.058 | 0.136 | -0.062 | 0.121 | 0.101 | 0.137 | 0.07 | 0.131 |
| Raw | LASSO | 0.118 | 0.114 | 0.01 | 0.166 | 0.198 | 0.141 | 0.141 | 0.153 |
| deriv1 | LASSO | 0.041 | 0.137 | 0.01 | 0.149 | 0.113 | 0.155 | 0.096 | 0.161 |
| Raw | RandomForest | -0.027 | 0.097 | 0 | 0.109 | 0.058 | 0.152 | 0.053 | 0.12 |
| deriv1 | RandomForest | 0.055 | 0.132 | -0.017 | 0.102 | 0.112 | 0.136 | 0.081 | 0.135 |
| Raw | RKHS | 0.033 | 0.192 | 0.021 | 0.199 | -0.008 | 0.187 | 0.051 | 0.184 |
| deriv1 | RKHS | 0.115 | 0.205 | 0.018 | 0.201 | 0.001 | 0.184 | 0.137 | 0.216 |
| Raw | rrBLUP | 0.017 | 0.119 | -0.026 | 0.113 | 0.103 | 0.147 | 0.019 | 0.143 |
| deriv1 | rrBLUP | 0.068 | 0.145 | -0.013 | 0.117 | 0.145 | 0.136 | 0.107 | 0.132 |

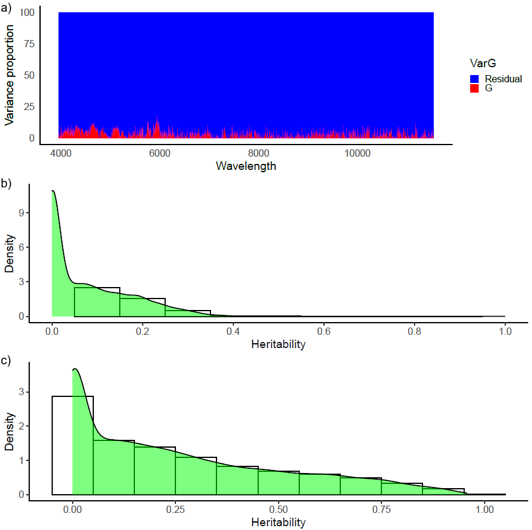

**Figure S3.** Genetic contribution to individual near-infrared wavelengths and metabolomic features. Panel a) shows variance proportions across NIR wavelengths, with genotypic and residual components in red and blue, respectively. Panels b) and c) show the distributions of feature-wise heritability estimates for NIR wavelengths and metabolomic features, respectively. NIR, near-infrared.

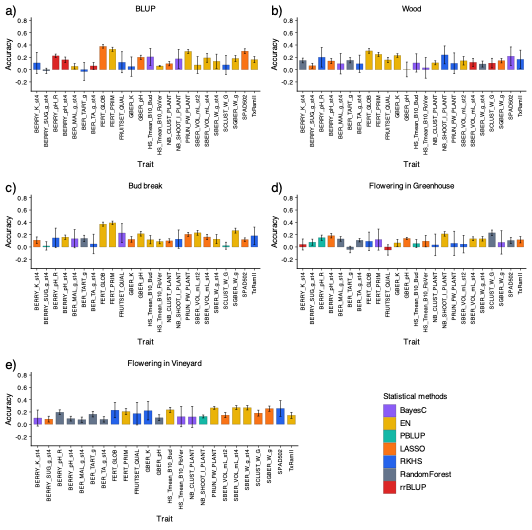

**Figure S4.** Trait-specific phenomic prediction models selected for each sampling category in population 50025. Panels a), b), c), d), and e) show combined-sample BLUPs, wood, vineyard leaves at budbreak, greenhouse leaves at flowering, and vineyard leaves at flowering, respectively. For each trait and sampling category, the model with the highest mean was retained. Colors denote models: purple, BayesC; yellow, elastic net; turquoise, PBLUP; orange, LASSO; blue, RKHS; grey, Random Forest; and red, rrBLUP. Error bars are standard deviations across 50 prediction replicates. BLUP, best linear unbiased prediction; LASSO, least absolute shrinkage and selection operator; PBLUP, phenomic relationship-matrix BLUP; RKHS, reproducing kernel Hilbert space; rrBLUP, ridge-regression BLUP.

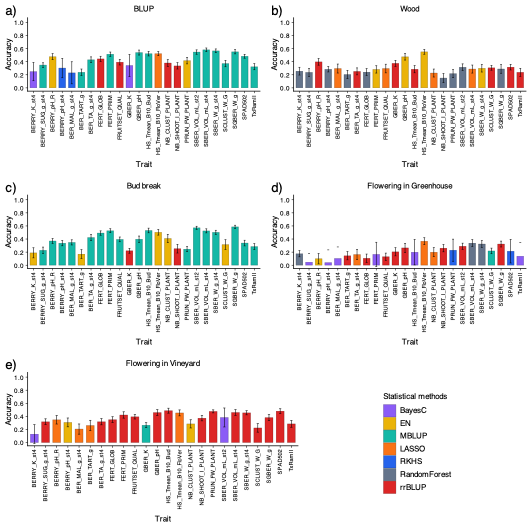

**Figure S5.** Trait-specific metabolomic prediction models selected for each sampling category in population 50025. Panels a), b), c), d), and e) show combined-sample BLUPs, wood, vineyard leaves at budbreak, greenhouse leaves at flowering, and vineyard leaves at flowering, respectively. For each trait and sampling category, the model with the highest mean was retained. Colors denote models: purple, BayesC; yellow, elastic net; turquoise, MBLUP; orange, LASSO; blue, RKHS; grey, Random Forest; and red, rrBLUP. Error bars are standard deviations across 50 prediction replicates. BLUP, best linear unbiased prediction; LASSO, least absolute shrinkage and selection operator; MBLUP, metabolomic relationship-matrix BLUP; RKHS, reproducing kernel Hilbert space; rrBLUP, ridge-regression BLUP.

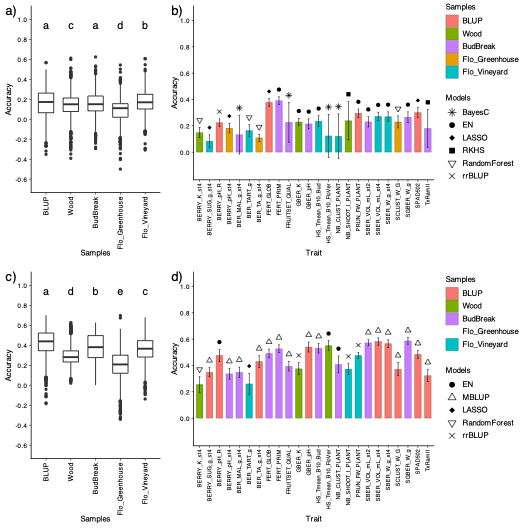

**Figure S6.** Effects of sampling category on phenomic and metabolomic prediction in population 50025. Panels a) and c) show predictive-ability distributions across traits for phenomic and metabolomic prediction, respectively; panels b) and d) show the highest-mean sampling-category and model combination for each trait. Letters above boxplots denote groups from Kruskal-Wallis tests followed by Dunn tests with Benjamini-Hochberg adjustment. Colors identify sampling categories: coral, combined-sample BLUPs; green, wood; purple, vineyard leaves at budbreak; orange, greenhouse leaves at flowering; and turquoise, vineyard leaves at flowering. Black symbols identify models: asterisks, BayesC; filled circles, Elastic Net; filled diamonds, LASSO; filled squares, RKHS; open downward-pointing triangles, Random Forest; diagonal crosses, rrBLUP; and open upward-pointing triangles, MBLUP. Error bars in panels b) and d) are standard deviations across 50 prediction replicates. BLUP, best linear unbiased prediction; LASSO, least absolute shrinkage and selection operator; MBLUP, metabolomic relationship-matrix BLUP; RKHS, reproducing kernel Hilbert space; rrBLUP, ridge-regression BLUP.

**Table S2.** Summary of leaf samples collected at flowering in the greenhouse and vineyard for each trait and omics dataset in the 50025 population. Mean and standard deviation of prediction accuracies for each combination of leaf samples collected at flowering (greenhouse and vineyard), trait, and omics dataset, using the highest-mean prediction models for each combination; SD: standard deviation.

| Traits | Metabolomic prediction | | | | Phenomic prediction | | | |
| --- | --- | --- | --- | --- | --- | --- | --- | --- |
|  | Greenhouse | | Vineyard | | Greenhouse | | Vineyard | |
|  | Mean | SD | Mean | SD | Mean | SD | Mean | SD |
| BERRY_K_st4 | 0.175 | 0.048 | 0.1 | 0.041 | 0.124 | 0.226 | 0.245 | 0.056 |
| BERRY_SUG_g_st4 | 0.041 | 0.054 | 0.318 | 0.044 | 0.19 | 0.074 | 0.054 | 0.131 |
| BERRY_pH_R | 0.1 | 0.079 | 0.348 | 0.065 | 0.069 | 0.032 | 0.046 | 0.046 |
| BERRY_pH_st4 | 0.144 | 0.2 | 0.307 | 0.068 | 0.126 | 0.173 | 0.114 | 0.143 |
| BER_MAL_g_st4 | 0.095 | 0.074 | 0.206 | 0.078 | 0.038 | 0.098 | 0.09 | 0.049 |
| BER_TART_g | 0.144 | 0.063 | 0.26 | 0.08 | -0.018 | 0.075 | 0.043 | 0.068 |
| BER_TA_g_st4 | 0.163 | 0.078 | 0.318 | 0.048 | 0.097 | 0.086 | -0.023 | 0.055 |
| FERT_GLOB | 0.106 | 0.064 | 0.35 | 0.044 | 0.154 | 0.16 | 0.189 | 0.051 |
| FERT_PRIM | 0.138 | 0.058 | 0.421 | 0.048 | 0.202 | 0.076 | 0.152 | 0.052 |
| FRUITSET_QUAL | 0.339 | 0.162 | 0.394 | 0.034 | 0.007 | 0.152 | 0.222 | 0.062 |
| GBER_K | 0.221 | 0.151 | 0.297 | 0.152 | -0.042 | 0.059 | 0.206 | 0.043 |
| GBER_pH | 0.265 | 0.07 | 0.458 | 0.043 | 0.026 | 0.167 | 0.112 | 0.054 |
| HS_Tmeam_B10_Bud | 0.189 | 0.049 | 0.487 | 0.038 | -0.033 | 0.059 | 0.334 | 0.038 |
| HS_Tmean_B10_FloVer | 0.367 | 0.053 | 0.453 | 0.046 | 0.108 | 0.165 | 0.159 | 0.059 |
| NB_CLUST_PLANT | 0.199 | 0.068 | 0.285 | 0.062 | 0.08 | 0.201 | 0.124 | 0.066 |
| NB_SHOOT_I_PLANT | 0.261 | 0.056 | 0.373 | 0.042 | 0.183 | 0.081 | 0.039 | 0.068 |
| PRUN_FW_PLANT | 0.226 | 0.051 | 0.481 | 0.111 | 0.077 | 0.054 | 0.256 | 0.031 |
| SBER_VOL_mL_st2 | 0.29 | 0.047 | 0.36 | 0.036 | 0.06 | 0.08 | 0.183 | 0.071 |
| SBER_VOL_mL_st4 | 0.338 | 0.052 | 0.46 | 0.043 | 0.233 | 0.144 | 0.179 | 0.037 |
| SBER_W_g_st4 | 0.324 | 0.05 | 0.454 | 0.031 | 0.214 | 0.146 | 0.178 | 0.046 |
| SCLUST_W_G | 0.217 | 0.039 | 0.223 | 0.064 | 0.298 | 0.059 | 0.043 | 0.06 |
| SGBER_W_g | 0.324 | 0.045 | 0.382 | 0.046 | 0.008 | 0.069 | 0.192 | 0.059 |
| SPAD502 | 0.206 | 0.054 | 0.48 | 0.036 | 0.085 | 0.046 | 0.264 | 0.042 |
| TxRamII | 0.12 | 0.057 | 0.286 | 0.051 | 0.035 | 0.056 | 0.047 | 0.007 |

**Table S3.** Summary of prediction accuracies across omics datasets and traits for the 50025 population. Mean and standard deviation of prediction accuracies for each omics dataset and trait, using the highest-mean prediction models and optimal samples for phenomic and metabolomic predictions.

|  | Genomic Prediction | | | Phenomic prediction | | | | Metabolomic prediction | | | |
| --- | --- | --- | --- | --- | --- | --- | --- | --- | --- | --- | --- |
| Traits | Mean | Standard deviation | Statistical models | Mean | Standard deviation | Samples | Statistical models | Mean | Standard deviation | Samples | Statistical models |
| BERRY_K_st4 | 0.326 | 0.076 | EN | 0.148 | 0.039 | Wood | RandomForest | 0.256 | 0.060 | Wood | RandomForest |
| BERRY_SUG_g_st4 | 0.548 | 0.043 | EN | 0.084 | 0.048 | Flowering Vineyard | LASSO | 0.349 | 0.034 | BLUP | MBLUP |
| BERRY_pH_R | 0.557 | 0.036 | rrBLUP | 0.224 | 0.029 | BLUP | rrBLUP | 0.477 | 0.046 | BLUP | EN |
| BERRY_pH_st4 | 0.308 | 0.036 | GBLUP | 0.182 | 0.037 | Flowering Greenhouse | LASSO | 0.337 | 0.040 | BudBreak | MBLUP |
| BER_MAL_g_st4 | 0.354 | 0.044 | rrBLUP | 0.133 | 0.148 | BudBreak | BayesC | 0.348 | 0.039 | BudBreak | MBLUP |
| BER_TART_g | 0.307 | 0.062 | EN | 0.164 | 0.045 | Flowering Vineyard | RandomForest | 0.260 | 0.080 | Flowering Vineyard | LASSO |
| BER_TA_g_st4 | 0.445 | 0.033 | rrBLUP | 0.109 | 0.027 | Flowering Greenhouse | RandomForest | 0.430 | 0.044 | BLUP | MBLUP |
| FERT_GLOB | 0.481 | 0.034 | rrBLUP | 0.378 | 0.028 | BLUP | LASSO | 0.491 | 0.033 | BudBreak | MBLUP |
| FERT_PRIM | 0.550 | 0.026 | rrBLUP | 0.391 | 0.029 | BudBreak | EN | 0.527 | 0.030 | BudBreak | MBLUP |
| FRUITSET_QUAL | 0.393 | 0.038 | GBLUP | 0.226 | 0.151 | BudBreak | BayesC | 0.394 | 0.036 | BudBreak | MBLUP |
| GBER_K | 0.379 | 0.043 | GBLUP | 0.229 | 0.028 | Wood | EN | 0.375 | 0.046 | Wood | rrBLUP |
| GBER_pH | 0.582 | 0.031 | GBLUP | 0.215 | 0.038 | BudBreak | EN | 0.540 | 0.038 | BLUP | MBLUP |
| HS_Tmeam_B10_Bud | 0.659 | 0.022 | rrBLUP | 0.235 | 0.041 | Flowering Vineyard | EN | 0.531 | 0.036 | BudBreak | MBLUP |
| HS_Tmean_B10_FloVer | 0.827 | 0.017 | EN | 0.123 | 0.164 | Flowering Vineyard | BayesC | 0.551 | 0.036 | Wood | EN |
| NB_CLUST_PLANT | 0.475 | 0.127 | BayesC | 0.120 | 0.171 | Flowering Vineyard | BayesC | 0.408 | 0.064 | BudBreak | EN |
| NB_SHOOT_I_PLANT | 0.278 | 0.134 | RKHS | 0.239 | 0.146 | Wood | RKHS | 0.373 | 0.042 | Flowering Vineyard | rrBLUP |
| PRUN_FW_PLANT | 0.344 | 0.046 | GBLUP | 0.296 | 0.031 | BLUP | EN | 0.476 | 0.023 | Flowering Vineyard | rrBLUP |
| SBER_VOL_mL_st2 | 0.688 | 0.091 | BayesC | 0.230 | 0.039 | BudBreak | EN | 0.572 | 0.024 | BudBreak | MBLUP |
| SBER_VOL_mL_st4 | 0.680 | 0.023 | GBLUP | 0.271 | 0.032 | Flowering Vineyard | EN | 0.582 | 0.028 | BLUP | MBLUP |
| SBER_W_g_st4 | 0.702 | 0.022 | GBLUP | 0.269 | 0.036 | Flowering Vineyard | EN | 0.566 | 0.029 | BLUP | MBLUP |
| SCLUST_W_G | 0.384 | 0.044 | rrBLUP | 0.230 | 0.045 | Flowering Greenhouse | RandomForest | 0.372 | 0.049 | BLUP | MBLUP |
| SGBER_W_g | 0.715 | 0.070 | BayesC | 0.266 | 0.040 | BudBreak | EN | 0.586 | 0.025 | BudBreak | MBLUP |
| SPAD502 | 0.420 | 0.051 | GBLUP | 0.301 | 0.037 | BLUP | LASSO | 0.484 | 0.030 | BLUP | MBLUP |
| TxRamII | 0.493 | 0.037 | rrBLUP | 0.181 | 0.143 | BudBreak | RKHS | 0.323 | 0.046 | BLUP | MBLUP |

**Table S4.** Summary of prediction accuracies across omics datasets and traits for the 50025 population. Mean and standard deviation of prediction accuracies for each combination of omics dataset and trait, based on the highest-mean prediction models and optimal samples for phenomic and metabolomic predictions.

| Trait | Genomic prediction | | Metabolomic prediction | | Phenomic prediction | | Genomic and metabolomic prediction | | Genomic and phenomic prediction | | Phenomic and metabolomic prediction | | Genomic, phenomic and metabolomic prediction | |
| --- | --- | --- | --- | --- | --- | --- | --- | --- | --- | --- | --- | --- | --- | --- |
|  | Mean | Standard deviation | Mean | Standard deviation | Mean | Standard deviation | Mean | Standard deviation | Mean | Standard deviation | Mean | Standard deviation | Mean | Standard deviation |
| BERRY_K_st4 | 0.326 | 0.076 | 0.256 | 0.060 | 0.148 | 0.039 | 0.302 | 0.113 | 0.332 | 0.171 | 0.259 | 0.133 | 0.301 | 0.135 |
| BERRY_SUG_g_st4 | 0.548 | 0.043 | 0.349 | 0.034 | 0.084 | 0.048 | 0.488 | 0.117 | 0.456 | 0.125 | 0.288 | 0.153 | 0.477 | 0.112 |
| BERRY_pH_R | 0.557 | 0.036 | 0.477 | 0.046 | 0.224 | 0.029 | 0.563 | 0.134 | 0.580 | 0.126 | 0.405 | 0.158 | 0.551 | 0.155 |
| BERRY_pH_st4 | 0.308 | 0.036 | 0.337 | 0.040 | 0.182 | 0.037 | 0.369 | 0.143 | 0.301 | 0.155 | 0.265 | 0.134 | 0.327 | 0.142 |
| BER_MAL_g_st4 | 0.354 | 0.044 | 0.348 | 0.039 | 0.133 | 0.148 | 0.375 | 0.136 | 0.334 | 0.140 | 0.267 | 0.182 | 0.307 | 0.193 |
| BER_TART_g | 0.307 | 0.062 | 0.260 | 0.080 | 0.164 | 0.045 | 0.256 | 0.162 | 0.185 | 0.137 | 0.225 | 0.167 | 0.255 | 0.179 |
| BER_TA_g_st4 | 0.445 | 0.033 | 0.430 | 0.044 | 0.109 | 0.027 | 0.417 | 0.112 | 0.422 | 0.150 | 0.289 | 0.163 | 0.389 | 0.134 |
| FERT_GLOB | 0.481 | 0.034 | 0.491 | 0.033 | 0.378 | 0.028 | 0.526 | 0.131 | 0.529 | 0.127 | 0.390 | 0.160 | 0.517 | 0.140 |
| FERT_PRIM | 0.550 | 0.026 | 0.527 | 0.030 | 0.391 | 0.029 | 0.552 | 0.124 | 0.565 | 0.116 | 0.425 | 0.147 | 0.555 | 0.132 |
| FRUITSET_QUAL | 0.393 | 0.038 | 0.394 | 0.036 | 0.226 | 0.151 | 0.441 | 0.142 | 0.390 | 0.163 | 0.353 | 0.149 | 0.423 | 0.135 |
| GBER_K | 0.379 | 0.043 | 0.375 | 0.046 | 0.229 | 0.028 | 0.411 | 0.147 | 0.434 | 0.126 | 0.296 | 0.180 | 0.419 | 0.142 |
| GBER_pH | 0.582 | 0.031 | 0.540 | 0.038 | 0.215 | 0.038 | 0.570 | 0.132 | 0.582 | 0.120 | 0.439 | 0.138 | 0.574 | 0.128 |
| HS_Tmeam_B10_Bud | 0.659 | 0.022 | 0.531 | 0.036 | 0.235 | 0.041 | 0.690 | 0.085 | 0.654 | 0.103 | 0.458 | 0.110 | 0.669 | 0.089 |
| HS_Tmean_B10_FloVer | 0.827 | 0.017 | 0.551 | 0.036 | 0.123 | 0.164 | 0.659 | 0.162 | 0.703 | 0.083 | 0.249 | 0.149 | 0.614 | 0.093 |
| NB_CLUST_PLANT | 0.475 | 0.127 | 0.408 | 0.064 | 0.120 | 0.171 | 0.472 | 0.135 | 0.457 | 0.132 | 0.294 | 0.158 | 0.435 | 0.141 |
| NB_SHOOT_I_PLANT | 0.278 | 0.134 | 0.373 | 0.042 | 0.239 | 0.146 | 0.305 | 0.152 | 0.317 | 0.141 | 0.308 | 0.140 | 0.320 | 0.164 |
| PRUN_FW_PLANT | 0.344 | 0.046 | 0.476 | 0.023 | 0.296 | 0.031 | 0.462 | 0.126 | 0.358 | 0.144 | 0.416 | 0.149 | 0.461 | 0.130 |
| SBER_VOL_mL_st2 | 0.688 | 0.091 | 0.572 | 0.024 | 0.230 | 0.039 | 0.676 | 0.109 | 0.681 | 0.082 | 0.442 | 0.141 | 0.661 | 0.119 |
| SBER_VOL_mL_st4 | 0.680 | 0.023 | 0.582 | 0.028 | 0.271 | 0.032 | 0.647 | 0.110 | 0.652 | 0.109 | 0.498 | 0.129 | 0.625 | 0.088 |
| SBER_W_g_st4 | 0.702 | 0.022 | 0.566 | 0.029 | 0.269 | 0.036 | 0.668 | 0.095 | 0.673 | 0.100 | 0.489 | 0.126 | 0.636 | 0.106 |
| SCLUST_W_G | 0.384 | 0.044 | 0.372 | 0.049 | 0.230 | 0.045 | 0.397 | 0.136 | 0.361 | 0.134 | 0.299 | 0.132 | 0.384 | 0.121 |
| SGBER_W_g | 0.715 | 0.070 | 0.586 | 0.025 | 0.266 | 0.040 | 0.678 | 0.077 | 0.679 | 0.095 | 0.439 | 0.124 | 0.675 | 0.095 |
| SPAD502 | 0.420 | 0.051 | 0.484 | 0.030 | 0.301 | 0.037 | 0.499 | 0.142 | 0.452 | 0.156 | 0.483 | 0.136 | 0.509 | 0.128 |
| TxRamII | 0.493 | 0.037 | 0.323 | 0.046 | 0.181 | 0.143 | 0.438 | 0.123 | 0.493 | 0.112 | 0.259 | 0.136 | 0.428 | 0.119 |

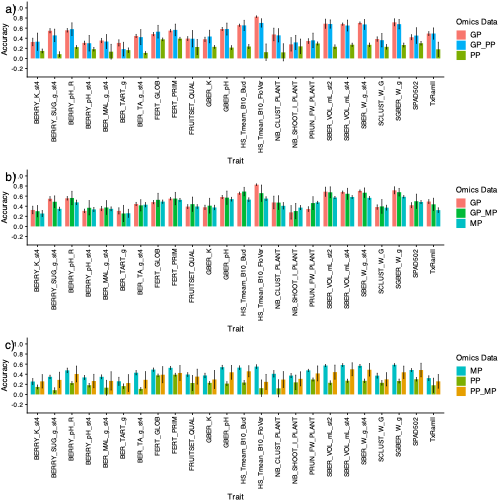

**Figure S7.** Effects of pairwise omics integration on prediction in population 50025. Panel a) compares genomic, phenomic, and combined genomic-phenomic prediction; panel b) compares genomic, metabolomic, and combined genomic-metabolomic prediction; panel c) compares phenomic, metabolomic, and combined phenomic-metabolomic prediction. For each trait and configuration, the model and, where applicable, sampling category with the highest mean were retained. Colors denote data layers: coral, GP; olive green, PP; turquoise, MP; blue, GP_PP; green, GP_MP; and orange, PP_MP. Error bars are standard deviations across 50 prediction replicates. GP, genomic prediction; MP, metabolomic prediction; PP, phenomic prediction.

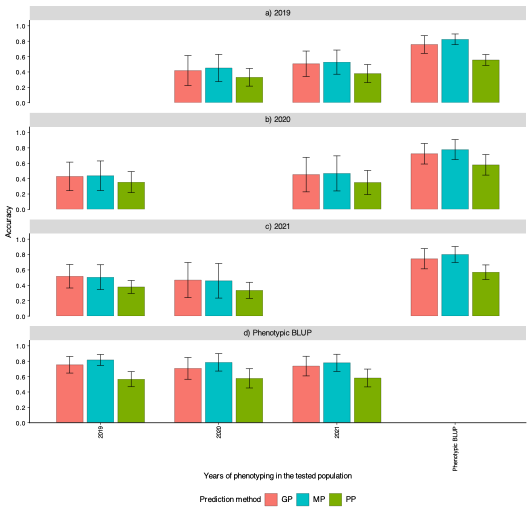

**Figure S8.** Influence of phenotyping year on prediction in population 50025.

Panel a) shows models trained with 2019 phenotypes and tested against 2020, 2021, or multi-year phenotypic BLUPs. Panels b) and c) show the corresponding analyses using 2020 and 2021 phenotypes for training, respectively. Panel d) shows models trained with phenotypic BLUPs and tested against each individual year. Training and testing used phenotypes from the same genotype set rather than disjoint genotype subsets. Bars and error bars represent means and standard deviations across traits after retaining the highest-mean model and, where applicable, sampling category. Coral, turquoise, and olive-green bars denote genomic, metabolomic, and phenomic prediction, respectively. BLUP, best linear unbiased prediction; GP, genomic prediction; MP, metabolomic prediction; PP, phenomic prediction.

**Table S5.** Summary of prediction accuracies across phenotyping years and omics datasets for the 50025 population. Mean and standard deviation of prediction accuracies for each omics dataset and combination of phenotyping years used for training and test populations, based on the highest-mean prediction models and optimal samples for phenomic and metabolomic predictions. BLUP refers to phenotypic BLUP.

| Train | Test | Genomic prediction | | Metabolomic prediction | | Phenomic prediction | |
| --- | --- | --- | --- | --- | --- | --- | --- |
|  |  | Mean | Standard deviation | Mean | Standard deviation | Mean | Standard deviation |
| 2019 | 2020 | 0.418 | 0.194 | 0.452 | 0.177 | 0.331 | 0.115 |
| 2019 | 2021 | 0.507 | 0.165 | 0.528 | 0.157 | 0.379 | 0.117 |
| 2019 | BLUP | 0.757 | 0.114 | 0.824 | 0.07 | 0.556 | 0.07 |
| 2020 | 2019 | 0.429 | 0.186 | 0.438 | 0.192 | 0.354 | 0.137 |
| 2020 | 2021 | 0.453 | 0.225 | 0.468 | 0.228 | 0.35 | 0.157 |
| 2020 | BLUP | 0.724 | 0.134 | 0.777 | 0.129 | 0.579 | 0.133 |
| 2021 | 2019 | 0.518 | 0.153 | 0.505 | 0.162 | 0.379 | 0.084 |
| 2021 | 2020 | 0.468 | 0.228 | 0.459 | 0.225 | 0.335 | 0.105 |
| 2021 | BLUP | 0.745 | 0.131 | 0.801 | 0.103 | 0.571 | 0.094 |
| BLUP | 2019 | 0.754 | 0.108 | 0.817 | 0.071 | 0.566 | 0.098 |
| BLUP | 2020 | 0.707 | 0.141 | 0.785 | 0.113 | 0.577 | 0.125 |
| BLUP | 2021 | 0.737 | 0.128 | 0.78 | 0.111 | 0.582 | 0.116 |

**Table S6.** Summary of trait prediction accuracies across populations using genomic and phenomic models. Mean and standard deviation of prediction accuracies for each omics dataset, cross-validation (CV) population, and trait, based on the highest-mean prediction models and optimal samples for phenomic prediction.

| Traits | Scenario | Genomic Prediction | | | Phenomic prediction | | |
| --- | --- | --- | --- | --- | --- | --- | --- |
|  |  | Mean | Standard deviation | Statistical models | Mean | Standard deviation | Statistical models |
| BER_MAL_g_st4 | CV-44910 | 0.286 | 0.041 | rrBLUP | 0.139 | 0.06 | EN |
| BER_MAL_g_st4 | CV-50025 | 0.386 | 0.043 | rrBLUP | 0.147 | 0.15 | BayesC |
| BER_TA_g_st4 | CV-44910 | 0.379 | 0.033 | rrBLUP | 0.216 | 0.04 | rrBLUP |
| BER_TA_g_st4 | CV-50025 | 0.499 | 0.026 | rrBLUP | 0.119 | 0.134 | RKHS |
| BER_TA_g_st4 | CV-RIxGW | 0.377 | 0.098 | BayesC | NA |  | NA |
| BER_TART_g | CV-44910 | 0.44 | 0.03 | GBLUP | 0.31 | 0.055 | EN |
| BER_TART_g | CV-50025 | 0.306 | 0.053 | EN | 0.108 | 0.071 | RandomForest |
| BERRY_K_st4 | CV-44910 | 0.332 | 0.041 | LASSO | 0.306 | 0.066 | EN |
| BERRY_K_st4 | CV-50025 | 0.328 | 0.063 | EN | 0.177 | 0.059 | LASSO |
| BERRY_K_st4 | CV-RIxGW | 0.146 | 0.044 | RandomForest | NA |  | NA |
| BERRY_pH_R | CV-44910 | 0.558 | 0.035 | rrBLUP | 0.147 | 0.074 | EN |
| BERRY_pH_R | CV-50025 | 0.645 | 0.026 | rrBLUP | 0.246 | 0.038 | rrBLUP |
| BERRY_pH_R | CV-RIxGW | 0.492 | 0.109 | BayesC | NA |  | NA |
| BERRY_pH_st4 | CV-44910 | 0.44 | 0.034 | GBLUP | 0.35 | 0.122 | BayesC |
| BERRY_pH_st4 | CV-50025 | 0.379 | 0.032 | rrBLUP | 0.168 | 0.048 | EN |
| BERRY_pH_st4 | CV-RIxGW | 0.455 | 0.083 | BayesC | NA |  | NA |
| BERRY_SUG_g_st4 | CV-44910 | 0.282 | 0.039 | rrBLUP | 0.128 | 0.105 | EN |
| BERRY_SUG_g_st4 | CV-50025 | 0.585 | 0.032 | EN | 0.119 | 0.064 | EN |
| BERRY_SUG_g_st4 | CV-RIxGW | 0.423 | 0.027 | GBLUP | NA |  | NA |
| FERT_GLOB | CV-44910 | 0.39 | 0.031 | rrBLUP | 0.145 | 0.132 | BayesC |
| FERT_GLOB | CV-50025 | 0.478 | 0.027 | rrBLUP | 0.324 | 0.065 | EN |
| FERT_GLOB | CV-RIxGW | 0.47 | 0.027 | rrBLUP | NA |  | NA |
| FERT_PRIM | CV-44910 | 0.42 | 0.021 | rrBLUP | 0.219 | 0.043 | LASSO |
| FERT_PRIM | CV-50025 | 0.592 | 0.024 | rrBLUP | 0.316 | 0.086 | EN |
| FERT_PRIM | CV-RIxGW | 0.408 | 0.081 | RKHS | NA |  | NA |
| FRUITSET_QUAL | CV-44910 | 0.208 | 0.035 | rrBLUP | 0.013 | 0.052 | GBLUP |
| FRUITSET_QUAL | CV-50025 | 0.364 | 0.04 | rrBLUP | 0.188 | 0.109 | LASSO |
| GBER_K | CV-44910 | 0.171 | 0.062 | RandomForest | 0.072 | 0.052 | GBLUP |
| GBER_K | CV-50025 | 0.404 | 0.036 | rrBLUP | 0.229 | 0.056 | EN |
| GBER_pH | CV-44910 | 0.555 | 0.024 | rrBLUP | 0.396 | 0.105 | BayesC |
| GBER_pH | CV-50025 | 0.639 | 0.023 | GBLUP | 0.189 | 0.036 | LASSO |
| GBER_pH | CV-RIxGW | 0.392 | 0.03 | GBLUP | NA |  | NA |
| HS_Tmeam_B10_Bud | CV-44910 | 0.594 | 0.021 | GBLUP | 0.27 | 0.075 | EN |
| HS_Tmeam_B10_Bud | CV-50025 | 0.631 | 0.026 | rrBLUP | 0.239 | 0.047 | LASSO |
| HS_Tmean_B10_FloVer | CV-44910 | 0.608 | 0.026 | rrBLUP | 0.13 | 0.117 | BayesC |
| HS_Tmean_B10_FloVer | CV-50025 | 0.822 | 0.013 | EN | 0.094 | 0.051 | LASSO |
| NB_CLUST_PLANT | CV-44910 | 0.134 | 0.063 | EN | 0.173 | 0.1 | LASSO |
| NB_CLUST_PLANT | CV-50025 | 0.475 | 0.035 | rrBLUP | 0.155 | 0.188 | BayesC |
| NB_SHOOT_I_PLANT | CV-44910 | 0.107 | 0.044 | rrBLUP | 0.099 | 0.033 | LASSO |
| NB_SHOOT_I_PLANT | CV-50025 | 0.186 | 0.042 | rrBLUP | 0.228 | 0.051 | LASSO |
| NB_SHOOT_I_PLANT | CV-RIxGW | 0.283 | 0.031 | GBLUP | NA |  | NA |
| PRUN_FW_PLANT | CV-44910 | NA |  | NA | NA |  | NA |
| PRUN_FW_PLANT | CV-50025 | 0.364 | 0.042 | GBLUP | NA |  | NA |
| PRUN_FW_PLANT | CV-RIxGW | 0.469 | 0.026 | rrBLUP | NA |  | NA |
| SBER_VOL_mL_st2 | CV-44910 | 0.521 | 0.031 | rrBLUP | 0.22 | 0.042 | EN |
| SBER_VOL_mL_st2 | CV-50025 | 0.712 | 0.02 | rrBLUP | 0.217 | 0.052 | LASSO |
| SBER_VOL_mL_st2 | CV-RIxGW | 0.461 | 0.115 | RKHS | NA |  | NA |
| SBER_VOL_mL_st4 | CV-44910 | 0.505 | 0.029 | rrBLUP | 0.203 | 0.045 | EN |
| SBER_VOL_mL_st4 | CV-50025 | 0.734 | 0.025 | rrBLUP | 0.199 | 0.057 | LASSO |
| SBER_VOL_mL_st4 | CV-RIxGW | 0.609 | 0.073 | BayesC | NA |  | NA |
| SBER_W_g_st4 | CV-44910 | 0.555 | 0.023 | rrBLUP | 0.233 | 0.043 | EN |
| SBER_W_g_st4 | CV-50025 | 0.754 | 0.021 | rrBLUP | 0.165 | 0.141 | BayesC |
| SBER_W_g_st4 | CV-RIxGW | 0.607 | 0.075 | BayesC | NA |  | NA |
| SCLUST_W_G | CV-44910 | 0.18 | 0.048 | GBLUP | 0.089 | 0.157 | RKHS |
| SCLUST_W_G | CV-50025 | 0.439 | 0.03 | rrBLUP | 0.174 | 0.137 | RKHS |
| SGBER_W_g | CV-44910 | 0.518 | 0.03 | rrBLUP | 0.232 | 0.044 | LASSO |
| SGBER_W_g | CV-50025 | 0.716 | 0.023 | rrBLUP | 0.222 | 0.052 | EN |
| SGBER_W_g | CV-RIxGW | 0.46 | 0.115 | RKHS | NA |  | NA |
| SPAD502 | CV-44910 | 0.312 | 0.036 | RandomForest | 0.091 | 0.049 | RandomForest |
| SPAD502 | CV-50025 | 0.496 | 0.03 | GBLUP | 0.206 | 0.024 | rrBLUP |
| TxRamII | CV-44910 | 0.166 | 0.039 | RandomForest | 0.087 | 0.059 | LASSO |
| TxRamII | CV-50025 | 0.546 | 0.026 | rrBLUP | 0.122 | 0.048 | RandomForest |

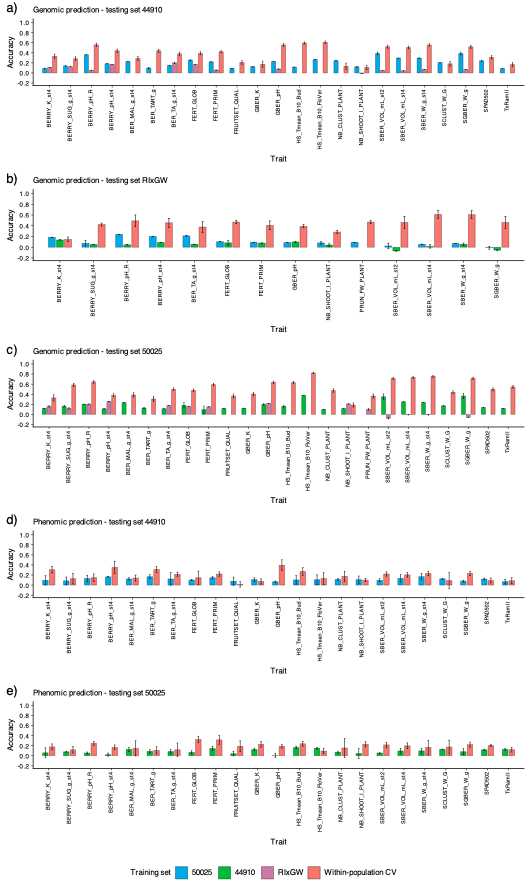

**Figure S9.** Genomic and phenomic prediction across training and testing populations.

Panel a) shows genomic prediction with population 44910 as the testing set; panel b) genomic prediction with RIxGW as the testing set; panel c) genomic prediction with population 50025 as the testing set; panel d) phenomic prediction with population 44910 as the testing set; and panel e) phenomic prediction with population 50025 as the testing set. Scenario direction is from the training population to the testing population. For each trait, the highest-mean eligible model and, for phenomic prediction, sampling category were retained. Blue, green, and magenta bars denote training with populations 50025, 44910, and RIxGW, respectively; coral bars denote within-population cross-validation. Error bars are standard deviations across 50 prediction runs where applicable; deterministic across-population fits may have a standard deviation of zero. CV, cross-validation.

**Table S7.** Summary of traits, phenotyping years, and BLUP availability. For berry maturation traits, stage 2 corresponds to measurements taken at the green berry stage, while stage 4 refers to measurements taken 380 degree days after véraison, representing mid-maturity.

| Code_EN | Trait | Year of phenotyping | BLUP several years |
| --- | --- | --- | --- |
| NB_SHOOT_I_PLANT | Number of primary branches | 2019-2020-2021 | 3 Years |
| HS_Tmeam_B10_Bud | Bud break date in degrees days with a base of 10 degrees Celsius | 2019-2020-2021 | 3 Years |
| HS_Tmean_B10_FloVer | Time between Flowering and Veraison in degrees day with a base of 10 degrees Celsius | 2019-2020-2021 | 3 Years |
| SPAD502 | Measure with SPAD502 | 2020-2021 | 2 Years |
| FRUITSET_QUAL | Quality of fruit set | 2020-2021 | 2 Years |
| SGBER_W_g | Weight of one berry at stage 2 | 2020-2021 | 2 Years |
| BERRY_pH_R | Berry pH at harvest | 2020-2021 | 2 Years |
| GBER_K | Berry potassium at stage 2 | 2020-2021 | 2 Years |
| GBER_pH | Berry pH at stage 2 | 2020-2021 | 2 Years |
| SBER_W_g_st4 | Weight of one berry at stage 4 | 2019-2020-2021 | 3 Years |
| SBER_VOL_mL_st2 | Volume of one berry at stage 2 | 2019-2020-2021 | 3 years |
| SBER_VOL_mL_st4 | Volume of one berry at stage 4 | 2019-2020-2021 | 3 years |
| BERRY_pH_st4 | Berry pH at stage 4 | 2019-2020-2021 | 3 Years |
| BERRY_SUG_g_st4 | Berry sugar content at stage 4 | 2019-2020-2021 | 3 Years |
| BER_TA_g_st4 | Total acidity in the berry at stage 4 | 2019-2020-2021 | 3 Years |
| BER_MAL_g_st4 | Berry malic acidity at stage 4 | 2019-2020-2021 | 3 Years |
| BER_TART_g | Berry tartaric acidity at stage 4 | 2019-2020-2021 | 3 Years |
| BERRY_K_st4 | Berry potassium at stage 4 | 2019-2020-2021 | 3 Years |
| NB_CLUST_PLANT | Harvest cluster number | 2019-2020-2021 | 3 Years |
| SCLUST_W_G | Harvest cluster weight | 2019-2020-2021 | 3 Years |
| FERT_PRIM | Primary fertility | 2019-2020-2021 | 3 Years |
| FERT_GLOB | Total fertility | 2019-2020-2021 | 3 Years |
| TxRamII | Secondary branches per plant rate | 2019-2020-2021 | 3 Years |
| PRUN_FW_PLANT | Weight of pruned wood | 2019-2020-2021 | 3 Years |

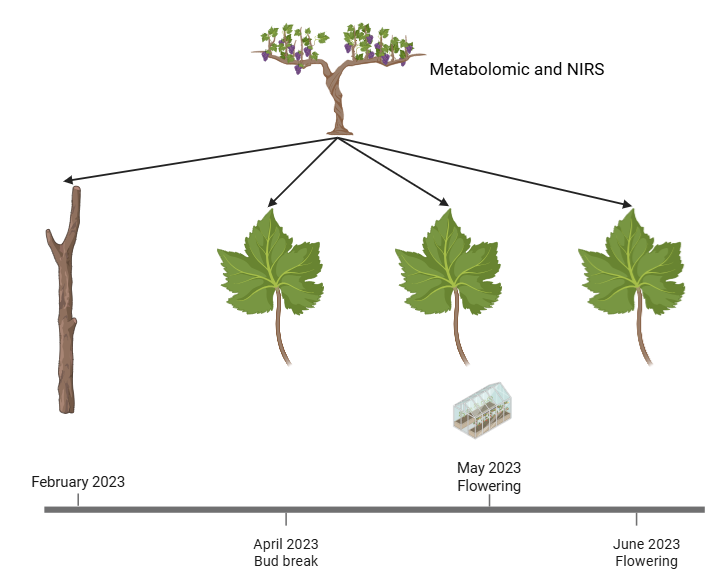

**Figure S10.** Sampling design for near-infrared and metabolomic profiling in population 50025.

Wood was collected in February 2023, vineyard leaves at budbreak in April 2023, greenhouse leaves at flowering in May 2023, and vineyard leaves at flowering in June 2023. The grapevine, wood, leaf, and greenhouse illustrations identify the source plant, sampled tissues, and controlled-environment collection; NIR, near-infrared. Created using biorender.com.
